# Synthetic essentiality of metabolic regulator PDHK1 in PTEN-deficient cells and cancers

**DOI:** 10.1101/441295

**Authors:** Nilanjana Chatterjee, Evangelos Pazarentzos, Gorjan Hrustanovic, Luping Lin, Erik Verschueren, Jeffrey R. Johnson, Matan Hofree, Jenny J. Yan, Victor Olivas, Billy W. Newton, John V. Dollen, Charles H. Earnshaw, Jennifer Flanagan, Elton Chan, Saurabh Asthana, Trey Ideker, Wei Wu, Manasi K. Mayekar, Junji Suzuki, Ben Barad, Yuriy Kirichok, James Fraser, William A. Weiss, Nevan J. Krogan, Asmin Tulpule, Amit J. Sabnis, Trever G. Bivona

## Abstract

PTEN is a tumor suppressor that is often inactivated in cancer and possesses both lipid and protein phosphatase activities. We report the metabolic regulator PDHK1 (pyruvate dehydrogenase kinase1) is a synthetic-essential gene in PTEN-deficient cancer and normal cells. The predominant mechanism of PDHK1 regulation and dependency is the PTEN protein phosphatase dephosphorylates NFκ;B activating protein (NKAP) and limits NFκB activation to suppress expression of PDHK1, a NFκB target gene. Loss of the PTEN protein phosphatase upregulates PDHK1 to drive aerobic glycolysis and induce PDHK1 cellular dependence. PTEN-deficient human tumors harbor increased PDHK1, which is a biomarker of decreased patient survival, establishing clinical relevance. This study uncovers a PTEN-regulated signaling pathway and reveals PDHK1 as a potential target in PTEN-deficient cancers.

**SIGNIFICANCE:** The tumor suppressor PTEN is widely inactivated in cancers and tumor syndromes. PTEN antagonizes PI3K/AKT signaling via its lipid phosphatase activity. The modest success of PI3K/AKT inhibition in PTEN-deficient cancer patients provides rationale for identifying other vulnerabilities in PTEN-deficient cancers to improve clinical outcomes. We show that PTEN-deficient cells are uniquely sensitive to PDHK1 inhibition. PTEN and PDHK1 co-suppression reduced colony formation and induced cell death *in vitro* and tumor regression *in vivo*. PDHK1 levels were high in PTEN-deficient patient tumors and associated with inferior patient survival, establishing clinical relevance. Our study identifies a PTEN-regulated signaling pathway linking the PTEN protein phosphatase to the metabolic regulator PDHK1 and provides a mechanistic basis for PDHK1 targeting in PTEN-deficient cancers.

## INTRODUCTION

*PTEN* was originally identified as a tumor suppressor gene and protein phosphatase (Li and Sun, 1997; Li et al., 1997; Steck et al., 1997) that is frequently genetically mutated, deleted, or epigenetically silenced in multiple cancer types and inactivated by germline genomic deletion in tumor syndromes and neurological disorders (Backman et al., 2001; Hollander et al., 2011; Kwon et al., 2006; Zhou and Parada, 2012). As a protein phosphatase, PTEN’s function was thought to directly suppress oncogenic tyrosine kinase signaling (Leslie et al., 2009). Subsequent biochemical characterization by several groups revealed that PTEN also possesses lipid phosphatase activity in addition to its dual-specificity protein phosphatase activity that favors phospho-tyrosine over phospho-serine/-threonine protein substrates (Maehama and Dixon, 1998; Maehama and Dixon, 1999; Myers et al., 1998; Myers et al., 1997). Importantly, as a lipid phosphatase PTEN directly antagonizes oncogenic PI3K/AKT/mTOR signaling through dephosphorylation of the lipid second messenger PIP3 (phosphatidylinositol-3,4,5 triphosphate) to PIP2 (phosphatidylinositol-4,5 bisphosphate), thereby regulating cellular metabolism, growth, and survival (Chalhoub and Baker, 2009; Lee et al., 1999; Maehama and Dixon, 1998; Myers et al., 1998; Stambolic et al., 1998).

Although the tumor suppressor function of PTEN is mainly attributed to its lipid phosphatase activity, PTEN can also function in a lipid phosphatase-independent (and also PI3K/AKT/mTOR-independent) manner through both protein phosphatase-dependent and lipid and protein phosphatase-independent (non-enzymatic) mechanisms (Davidson et al., 2010; Freeman et al., 2003; Gildea et al., 2004; Leslie et al., 2009; Maier et al., 1999; Planchon et al., 2008; Shen et al., 2007; Song et al., 2012). Consistent with the notion of PTEN lipid phosphatase-independent functions, PI3K and AKT inhibitors have shown modest clinical efficacy to date in most patients with PTEN-deficient cancers (Chandarlapaty et al., 2011; Ghosh et al., 2013; Rodon et al., 2013). Further, the existence of tumor-derived *PTEN Y138* mutants that selectively lack protein phosphatase activity, but retain lipid phosphatase activity suggests protein phosphatase-dependent cellular processes may contribute to PTEN function and tumor suppression (Davidson et al., 2010; Tibarewal et al., 2012). Recent studies also identified several signaling proteins as PTEN protein phosphatase substrates, including Rab7 (Shinde and Maddika, 2016), FAK (You et al., 2015), IRS1 (Shi et al., 2014), CREB (Gu et al., 2011), PDGF receptor (Mahimainathan and Choudhury, 2004) and PTK6 (Wozniak et al., 2017) and demonstrated that the protein phosphatase activity of PTEN can control important cellular processes including cell invasion (Tibarewal et al., 2012), migration (Dey et al., 2008; Leslie et al., 2007; Raftopoulou et al., 2004), cell-cycle progression (Hlobilkova et al., 2000), endocytic trafficking (Shinde and Maddika, 2016) and epithelial-to-mesenchymal transition (Leslie et al., 2007). Additionally, the protein phosphatase activity of PTEN may be important for its autoregulation (Zhang et al., 2012).

However, the signaling mechanisms by which the PTEN protein phosphatase functions, its direct substrates and effectors remain incompletely characterized, as does the role(s) of the PTEN protein phosphatase in normal cells and cancer. We investigated the molecular events regulated by the PTEN protein phosphatase in normal and cancer cells to provide insight into activity-specific PTEN functions and to identify unrecognized vulnerabilities that could be therapeutically exploited to improve the treatment of PTEN-deficient cancers.

## RESULTS

### PTEN deficiency upregulates PDHK1 expression in normal and cancer cells

Cancer cells offer a useful model system to investigate PTEN-related signaling and function, given that many cancers show loss of PTEN function through genetic or epigenetic mechanisms (Hollander et al., 2011). To identify uncharacterized molecular events through which PTEN functions in cells, we first investigated gene expression changes in PTEN-deficient human lung adenocarcinoma H1650 cells (Table S1), in comparison to the matched cell line into which wild-type (WT) PTEN was re-introduced. Stable PTEN re-expression in H1650 cells suppressed the levels of phosphorylated (p)-AKT (Figure 1A), as expected (Sos et al., 2009) and resulted in minimal growth effects (Figure S1A), consistent with prior work (Takeda et al., 2013).

**Figure 1.**
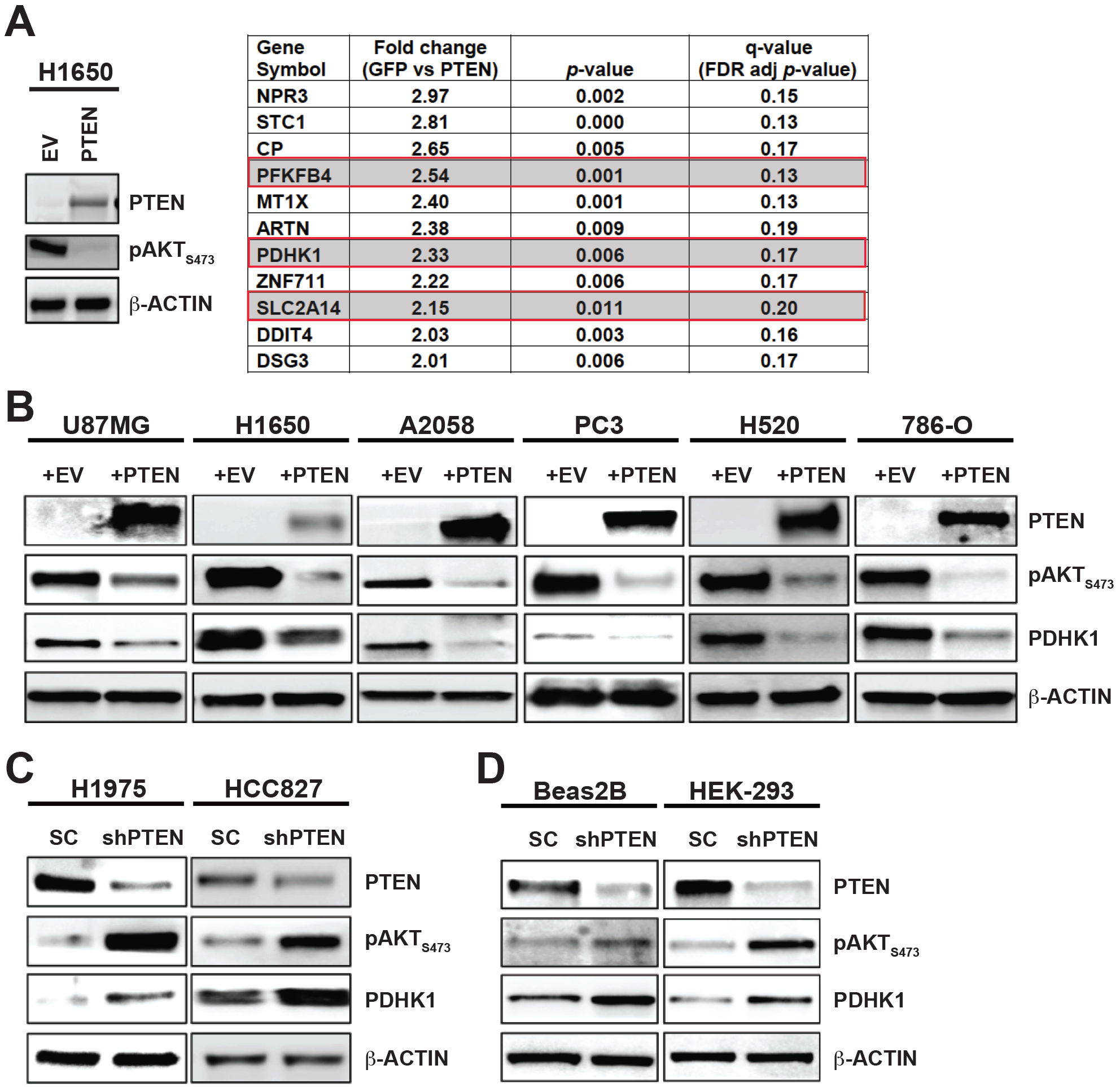
PTEN loss or inactivation upregulates PDHK1 in cancer and normal cells. (A) (Left) Western blots showing PTEN and phospho-AKT expression in PTEN-deficient H1650 cancer cell line stably expressing PTEN^WT^ or empty vector (EV). (Right) List of genes significantly upregulated (fold change > 2, multiple t-test \**p* < 0.05, FDR adjustedp-value or *q* < 0.2) in PTEN-deficient H1650-GFP cells compared to H1650 cells engineered to re-express PTEN stably by microarray analysis. Highlighted in red boxes are the top upregulated energy metabolism genes, including *PDHK1*, in H1650-GFP cells. (B) Western blots showing PTEN, phospho-AKT and PDHK1 expression in PTEN-deficient cancer cell lines stably expressing PTEN^WT^ or empty vector. (C-D) Same as (B) in PTEN-proficient cancer (C) or non-cancer (normal) (D) cell lines with stable *PTEN* knockdown. shPTEN, shRNA to PTEN and SC, scrambled control shRNA. See also Figures S1-S3 and Tables S1-S2.

By unbiased comparative gene expression profiling analysis, we identified 52 significantly differentially regulated genes (fold change > 2, \**p* < 0.05, *q* < 0.2) between the PTEN re-expressing and the parental PTEN-deficient H1650 cells (Figure 1A, and Table S2). Re-introduction of functional PTEN in H1650 cells resulted in statistically significant downregulation of 11 genes and upregulation of 41 genes. Interestingly, we found several energy metabolism genes including pyruvate dehydrogenase kinase 1 (PDHK1, gene name, *PDK1*) among the genes that showed statistically significant upregulation (>2-fold increase, \**p* < 0.05, *q* < 0.2) in the PTEN-deficient H1650 parental cells (Figure 1A, red boxes).

PDHK1 is a critical regulator of energy metabolism in both normal and cancer cells (Schulze and Downward, 2011) that has no known connection to PTEN, unlike the other top hit 6-phosphofructo-2-kinase/fructose-2,6-bisphosphatase 4 (PFKFB4) (Figure 1A, red boxes) which is regulated by AKT, a known downstream effector of PTEN (Chesney et al., 2014; Figueiredo et al., 2017; Houddane et al., 2017; Sun et al., 1999). PDHK1 and other PDHK isoenzymes (PDHK2-4) phosphorylate the E1 alpha subunit (PDHA1) of the pyruvate dehydrogenase complex (PDC) that catalyzes oxidative decarboxylation of pyruvate to acetyl-CoA in mitochondria (Patel and Roche, 1990; Popov et al., 1997; Teague et al., 1979). Phosphorylation of PDHA1 inhibits PDC activity and blocks pyruvate entry into the TCA cycle to uncouple glycolysis from the TCA cycle, thus contributing to the Warburg effect (Korotchkina and Patel, 2001; Linn et al., 1969; Vander Heiden et al., 2009; Warburg, 1956). Interestingly, analysis of a publicly-available dataset of an independently-conducted large-scale RNA interference screening project (Project Achilles, Broad Institute, http://www.broadinstitute.org/achilles) showed that 3 out of 4 PDHK1-targeted shRNAs were depleted in H1650 cells, in which we observed PDHK1 upregulation (Figure 1A). These findings suggest that PDHK1 expression may be essential for growth or survival in PTEN-deficient H1650 cells. Based on the collective data, we hypothesized that PDHK1 is an unrecognized PTEN effector through which PTEN regulates cellular energy metabolism and survival.

To investigate the potential role of PDHK1 as a PTEN effector, we assessed whether PDHK1 expression was regulated by PTEN more generally across a panel of paired cancer and normal cell lines that either lack or stably express PTEN (Table S1) (Aguissa-Toure and Li, 2012; Koul, 2008; Lee et al., 2014; Pourmand et al., 2007; Sos et al., 2009). Stable PTEN reexpression in PTEN-deficient cells (or *PTEN* knockdown in PTEN-proficient cells) resulted in minimal growth effects, as before (Figures S1A-S1G). We found PDHK1 expression was decreased by PTEN re-expression in multiple PTEN-deficient cancer cell types, such as lung adenocarcinoma (H1650), lung squamous cell carcinoma (H520), glioblastoma (U87MG), melanoma (A2058), prostate adenocarcinoma (PC3), and renal carcinoma cells (786-O) (Figures 1B, S2A, and Table S1). Conversely, PDHK1 expression was increased upon PTEN silencing in cancer cells and in normal (non-cancer) cells that are otherwise PTEN-proficient, such as multiple lung adenocarcinoma models (HCC827, H1975) and non-cancer cells (Beas2B, HEK293T) (Figures 1C, 1D, S2A, and Table S1). Together, these findings indicate an inverse relationship in which PTEN status (i.e. proficiency or deficiency) controls PDHK1 expression in normal and cancer cells. Further, unlike PDHK1, other PDHK isoforms including PDHK2, 3 and 4 exhibited lower expression in cancer cells consistent with earlier studies (Grassian et al., 2011) and were not significantly modulated by PTEN levels in these systems (Figure S2B), suggesting specificity in the link between PTEN and PDHK1. Altogether, these findings indicate that PTEN loss induces PDHK1 expression in normal and cancer cells, and suggest a specific mechanistic link between PTEN and PDHK1.

**Figure 2.**
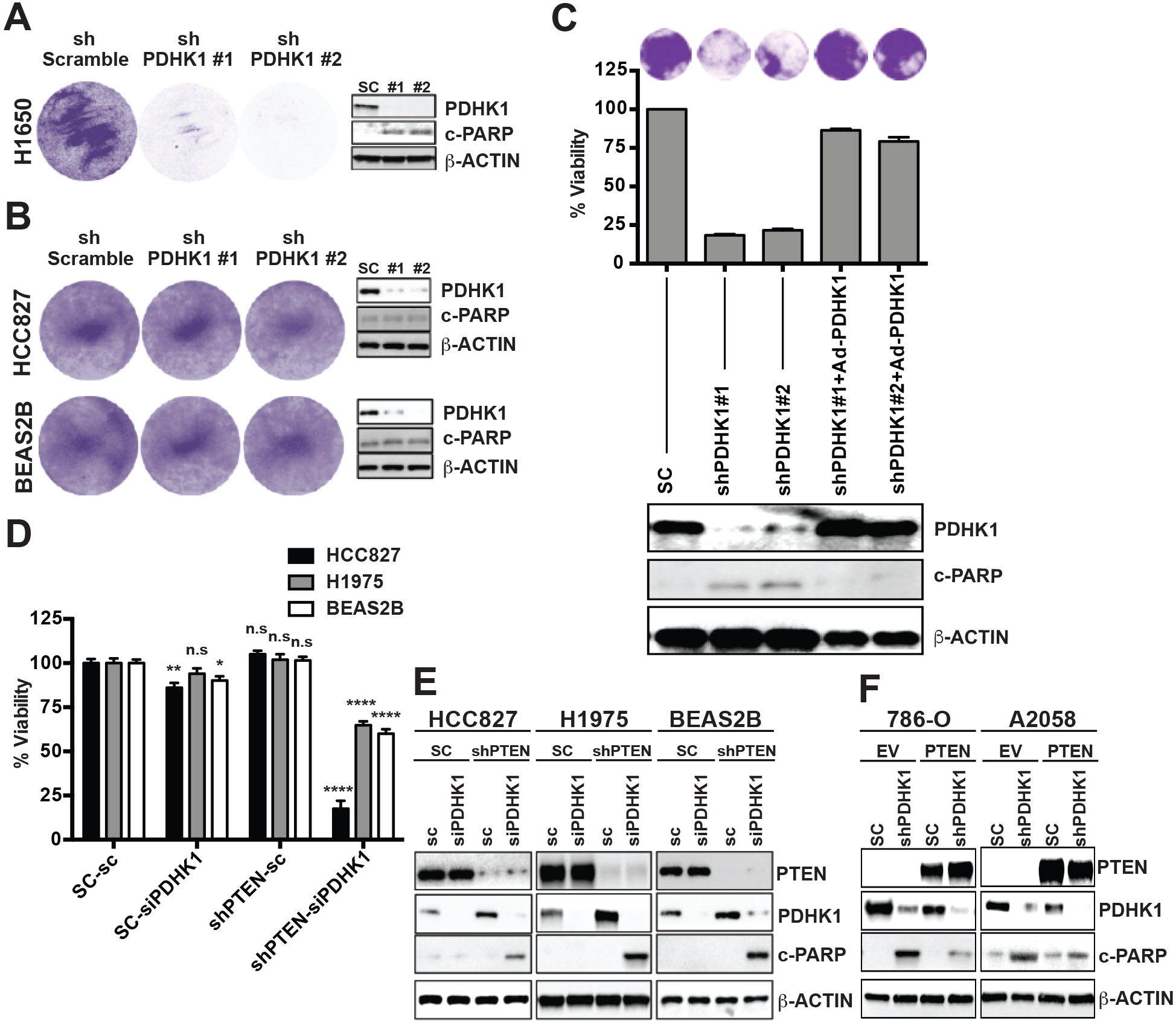
PTEN loss or inactivation induces cellular dependence on PDHK1 for survival. (A-B) Effects of stable *PDHK1* knockdown in PTEN-deficient H1650 cancer cell line (A) or PTEN-proficient cancer (HCC827) and normal (Beas2B) cell lines (B) on cell growth by crystal violet staining assay (left) and apoptosis induction as measured by cleaved PARP levels by immunoblot analysis (right) are shown. shPDHK1#1 and shPDHK1#2, shRNAs to PDHK1 and shScramble, scrambled control shRNA. (C) Effects of stable *PDHK1* knockdown and adenovirus-based shRNA resistant PDHK1 re-expression in PTEN-deficient H1650 cancer cell line on cell growth by crystal violet staining assay (top) or CellTiter-Glo luminescent assay (middle) and apoptosis induction as measured by cleaved PARP levels by immunoblot analysis (bottom) are shown. shPDHK1#1 and shPDHK1#2, shRNAs to PDHK1 and SC, scrambled control shRNA. (D-E) Effects of transient *PDHK1* knockdown in PTEN-proficient cancer and normal cell lines with or without stable *PTEN* knockdown on cell viability by crystal violet staining assay (D) and apoptosis induction as measured by cleaved PARP levels by immunoblotting (E) are shown. siPDHK1, PDHK1 specific small interfering RNAs and sc, scrambled control siRNA. shPTEN, shRNA to PTEN and SC, scrambled control shRNA. Data are shown as mean ± SEM (n = 4 replicates). \*\*\*\**p* < 0.0001; \*\**p* < 0.01; \**p* < 0.05; n.s, not significant compared to ‘scrambled control siRNA and shRNA expressing PTEN-proficient cells’ by Tukey’s multiple comparisons one-way ANOVA test. (F) Effects of stable *PDHK1* knockdown in PTEN-deficient cancer cell lines stably expressing PTEN or empty vector on apoptosis induction as measured by cleaved PARP levels by immunoblotting are shown. shPDHK1, shRNA to PDHK1 and sc, scrambled control shRNA. See also Figures S4-S6.

Using an independently-generated genetic dataset (Pathak et al., 2013), we corroborated these *in-vitro* findings *in-vivo* and in non-malignant tissue. Increased PDHK1 levels were present in normal murine lung epithelium in which PTEN was genetically-inactivated *in-vivo* compared to lung epithelium from PTEN^WT^ control mice (Figure S2C), suggesting physiological regulation of PDHK1 expression by PTEN *in-vivo*.

To establish the clinical relevance of our findings, we next investigated whether PTEN deficient human cancer specimens harbor increased PDHK1 levels. By immunohistochemistry (IHC) analysis in clinical specimens, we found that there was a significant inverse relationship between PDHK1 and PTEN expression levels in multiple human tumor types with frequent PTEN inactivation, including glioblastoma, prostate adenocarcinoma, and lung squamous cell carcinoma (Figures S3A-S3F). Additionally, we found a strong association between PTEN inactivation and increased PDHK1 expression in a pan-cancer analysis across 12 different tumor types in the Cancer Genome Atlas (TCGA) dataset (Figure S3G). Together, these data reinforced the PTEN/PDHK1 inverse relationship in normal and cancer cells that we observed in cell-based functional studies.

By analyzing the range of PDHK1 expression levels in these PTEN-deficient cancers, we also found that higher PDHK1 expression was a biomarker of decreased patient survival, suggesting that PDHK1 may contribute to PTEN’s tumor suppressor function (Figure S3H). Altogether, our data reveal that PTEN loss promotes PDHK1 expression in malignant and normal cells and tissues, both *in-vitro* and *in-vivo* and uncover increased PDHK1 levels in PTEN-deficient patient tumors as a biomarker of worse survival outcome.

### PTEN and PDHK1 co-suppression confers synthetic lethality in normal and cancer cells

We next investigated whether the PDHK1 upregulation induced by PTEN inactivation in normal and cancer cells is important for cell growth and survival. By suppressing PDHK1 using both genetic (shRNAs to knockdown *PDHK1*, Figures 2A, 2B, S4A and S4B) and validated pharmacologic (DCA or dichloroacetate (Kato et al., 2007; Stacpoole, 1989) approaches to inhibit PDHK1 (Figure S4C), we found that PDHK1 was essential for the survival of PTEN-deficient, but not PTEN-expressing, cancer and normal cells. We ruled out potential off-target effects of shRNA knockdown in this system, as the synthetic lethality conferred upon by PDHK1 co-suppression with PTEN loss in PTEN-deficient cells was rescued by re-expression of shRNA-resistant PDHK1 in these cells (Figure 2C).

Further, we confirmed that DCA treatment suppressed phosphorylation of the PDHK1 target PDHA1, as expected (Whitehouse et al., 1974), in PTEN-deficient cells (Figure S4D). While DCA may have off-target effects, the primary activity in the PTEN-deficient cells is likely via PDHK1 inhibition in these cells, as other known targets of DCA (e.g. PDHK2-4) were either not expressed or not PTEN-responsive in these systems (Figure S2B). Furthermore, PTEN status specifically dictated sensitivity to targeted PDHK1 inhibition, but not more generally to cytotoxic chemotherapies (Figure S4E).

To further establish that PTEN inactivation drives a functional PDHK1 dependence in cells, we employed the genetically-controlled system of paired cell lines that we engineered to stably lack or express PTEN. *PDHK1* knockdown significantly decreased survival in normal and cancer cells specifically in the context of PTEN co-suppression (Figure 2D). PDHK1 cosuppression with PTEN inactivation or loss was lethal in both cancer and normal cells, with biochemical evidence of enhanced apoptosis (measured by cleaved-PARP levels) in the models studied (Figures 2E and 2F). Similarly, PDHK1 inhibition with DCA decreased p-PDHA1 levels, as expected (Whitehouse et al., 1974), and increased apoptosis specifically in PTEN-deficient cells with increased PDHK1 (Figure S4F). Thus, PDHK1 is essential for survival in these PTEN-deficient cell systems.

We extended these findings by showing that PDHK1 inhibition by DCA treatment suppressed colony formation *in-vitro* specifically in PTEN-deficient lung adenocarcinoma cells (Figure S5A) and tumor growth *in-vivo* in PTEN-deficient melanoma xenograft models (Figure S5B). Overall, these data indicate that PDHK1 is conditionally essential for growth and survival specifically in PTEN-deficient cells and tumors.

### PDHK1 upregulation and activation upon PTEN loss promotes aerobic glycolysis

We further explored the functional consequences of PDHK1 upregulation induced by PTEN inactivation. PTEN and PDHK1 each can regulate cellular energy metabolism, but are not known to function together (Gottlob et al., 2001; Jang et al., 2013; Semenza, 2008). PDHK1 is a regulator of the metabolic switch from oxidative phosphorylation to aerobic glycolysis. PDHK1 activation can promote aerobic glycolysis, through which pyruvate is converted to lactate to produce ATP, by phosphorylating and inactivating PDHA1 and thereby blocking pyruvate entry into the TCA cycle (Patel et al., 2014). We therefore investigated the metabolic effects of PDHK1 upregulation in cells with PTEN deficiency.

First, we found that silencing of *PTEN* in cells induced L-lactate secretion, a hallmark of aerobic glycolysis (Figure S6A). Similarly, PTEN-deficient cells also exhibited increased L-lactate production in comparison to when these cells were grown in the presence of 2-DG (2-deoxyglucose, a glucose derivative that cannot undergo further glycolysis (Schulze and Harris, 2012; Zhao et al., 2013)) that suppresses lactate production independently (Figure S6B). Conversely, PDHK1 suppression by shRNA or by DCA treatment, or by stable PTEN reexpression in PTEN-deficient cell lines decreased L-lactate production (Figure S6B). These data suggest that PDHK1 upregulation and activation that occurs upon loss of PTEN promotes aerobic glycolysis in cells.

### PTEN protein phosphatase deficiency activates PDHK1 independent of PI3K/AKT and induces vulnerability to PDHK1 inhibition

Having identified a conditionally synthetic essential function of PDHK1 in cells lacking PTEN and the metabolic effects of PDHK1 activation in PTEN-deficient cells, we next investigated the mechanism by which PTEN regulates PDHK1 expression. We determined the requirement of the distinct lipid and protein phosphatase activities of PTEN in the regulation of PDHK1. We leveraged established cancer-derived PTEN mutants that abrogate either PTEN’s lipid phosphatase activity only (PTEN^G129E^) or its protein phosphatase activity only (PTEN^Y138L^) or both (PTEN^C124S^) (Davidson et al., 2010; Myers et al., 1998; Myers et al., 1997; Rodriguez-Escudero et al., 2011; Tibarewal et al., 2012). By stably expressing each of these PTEN mutants or PTEN^WT^ in a controlled system of PTEN-deficient cells, we found that the protein phosphatase, but not the lipid phosphatase, activity of PTEN regulates PDHK1 mRNA expression (Figure 3A). We confirmed that both mRNA and protein expression levels of PDHK1 were diminished upon expression of PTEN^WT^ or the PTEN mutant that retains the protein phosphatase activity only (PTEN^G129E^, lipid phosphatase mutant) in comparison to PTEN-deficient parental cells (Figures 3A and S7A). Further, PDHK1 protein levels were unaffected by expression of the PTEN mutant that lacks both phosphatase activities (PTEN^C124S^), suggesting that PDHK1 regulation by PTEN is phosphatase-activity dependent and not via other known non-enzymatic properties of PTEN (Freeman et al., 2003; Planchon et al., 2008; Shen et al., 2007) (Figure S7A).

**Figure 3.**
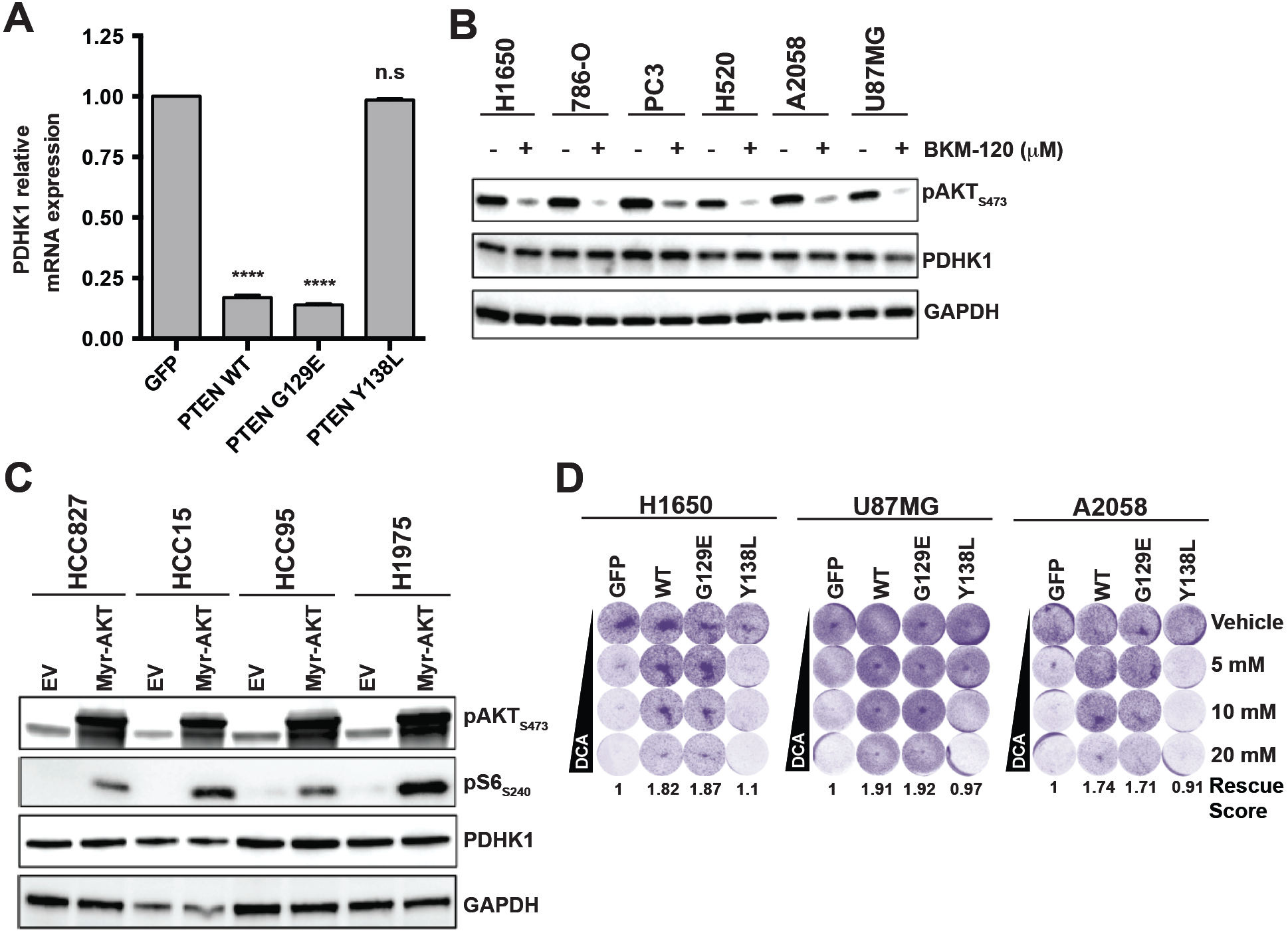
PTEN protein phosphatase and not lipid phosphatase represses PDHK1 independent of PI3K/AKT and loss of PTEN protein phosphatase renders PDHK1 essential for cell survival. (A) Quantitative real-time PCR analysis of PDHK1 mRNA expression in PTEN-deficient A2058 cancer cell line stably expressing PTEN^WT^ or PTEN^G129E^ or PTEN^Y138L^ or GFP. Data are shown as mean ± SD (n = 2 replicates). \*\*\*\**p* < 0.0001; n.s, not significant compared to ‘GFP expressing PTEN-deficient cells’ by Tukey’s multiple comparisons one-way ANOVA test. (B) Western blots showing phospho-AKT and PDHK1 expression in PTEN-deficient cancer cell lines in response to 1 μM BKM-120 (PI3-kinase inhibitor) or vehicle treatment for 24 hrs. (C) Western blots showing phospho-AKT, phospho-S6 (mTOR effector) and PDHK1 expression in PTEN-proficient cancer cell lines expressing empty vector (EV) or myristoylated-AKT (Myr-AKT) to constitutively activate AKT signaling. (D) Effects of pharmacologic inhibition of PDHK1 with DCA (dose response: 5, 10 and 20 mM) in PTEN-deficient stable cancer cell lines expressing PTEN^WT^ or PTEN^G129E^ or PTEN^Y138L^ or GFP on cell growth by crystal violet staining assay are shown, with quantification of cell viability in 20 mM DCA treatment relative to vehicle (water) treatment reported as rescue score (Methods). See also Figure S7.

As PDHK1 levels were unaffected by the presence or absence of the PTEN lipid phosphatase, we further investigated whether PDHK1 activation in PTEN-deficient cells was dependent upon PI3K/AKT signaling. Pharmacologic inhibition of PI3K or AKT in PTEN-deficient cells treated with BKM-120 (Burger et al., 2011) or MK2206 (Hirai et al., 2010), respectively, suppressed AKT activation (measured by phospho-AKT levels) and also expression of hexokinase 2 (HK2) (Figures S7B and S7C), which was upregulated in PTEN-deficient cells consistent with earlier studies (Wang et al., 2014). In contrast, the expression of PDHK1 was unaffected under these conditions of PI3K/AKT blockade (Figures 3B and S7B-S7D).

AKT is known to inactivate glycogen synthase kinase 3 α/β (GSK3α/β) through phosphorylation (Cross et al., 1995). Consistent with this, p-GSK3α/β (phospho-glycogen synthase kinase 3 α/β) levels were suppressed in PTEN-deficient cell lines by BKM-120 treatment, confirming the efficacy of the inhibitor; however, PDHK1 expression was not significantly affected (Figure S7D). Further, treatment with BKM-120 (Burger et al., 2011), and another PI3K inhibitor GDC0941 (Folkes et al., 2008), suppressed phosphorylation of ribosomal protein S6, which is downstream of both AKT and mTORC1, in 3 different PTEN-deficient cell lines tested and consistent with prior studies (Neshat et al., 2001), without decreasing PDHK1 levels (Figures S7D and S7E).

Conversely, ectopic expression of a constitutively active form of AKT (Myr-AKT) (Fulton et al., 1999; Sun et al., 2014), while increasing phospho-S6 levels as expected (Wittenberg et al., 2016), did not increase PDHK1 levels in PTEN-expressing cells (Figure 3C). Together, these findings indicate that PDHK1 is upregulated specifically by PTEN protein phosphatase inactivation, and in a PI3K/AKT-independent manner.

Additionally, we found that it was the protein phosphatase but not lipid phosphatase activity of PTEN that was required to rescue PTEN-deficient cells from the lethal effects of PDHK1 inhibition (Figure 3D). These collective data indicate that PDHK1 is conditionally essential for the survival specifically of PTEN protein phosphatase-deficient cells.

### PTEN protein phosphatase suppresses NFκB activation to repress PDHK1

We next investigated the mechanism by which PTEN protein phosphatase inactivation increases *PDHK1* gene (and thus protein) expression. Although PDHK1 expression can be upregulated by the transcription factor hypoxia-inducible factor 1 (HIF-1) (Kim et al., 2006), silencing the HIF-1α subunit of HIF-1 failed to suppress PDHK1 expression in a panel of PTEN-deficient cell lines, unlike the effects of HIF1α silencing in these cells on carbonic anhydrase 9 (CA9), which is known to be regulated by HIF-1 (Figure S8A) (Wykoff et al., 2000).

To understand the molecular basis of the transcriptional upregulation of PDHK1 that is induced by PTEN inactivation in cells, we sought to identify the transcription factor/s involved. We analyzed putative transcription factor binding sequences/motifs in, or upstream of the PDHK1 promoter by an *in-silico* promoter analysis. We identified a putative nuclear factor kappa-light chain-enhancer of activated B cells (NFκB) transcription factor (Gilmore, 2006) consensus DNA binding site (GGGRNNYYCC; R is purine, Y is pyrimidine and N is any base) (Martone et al., 2003) GGGACGCTCC at nucleotide position 173420479 in chromosome 2, at ~300 bp upstream of the TSS or transcriptional start site in the PDHK1 promoter, among other putative transcription factor consensus binding sequences (Figures 4A, S8B and Table S3). Based on these findings, we tested the hypothesis that PDHK1 is a transcriptional target of the transcription factor NFκB, which may directly upregulate PDHK1 mRNA expression upon PTEN inactivation.

**Figure 4.**
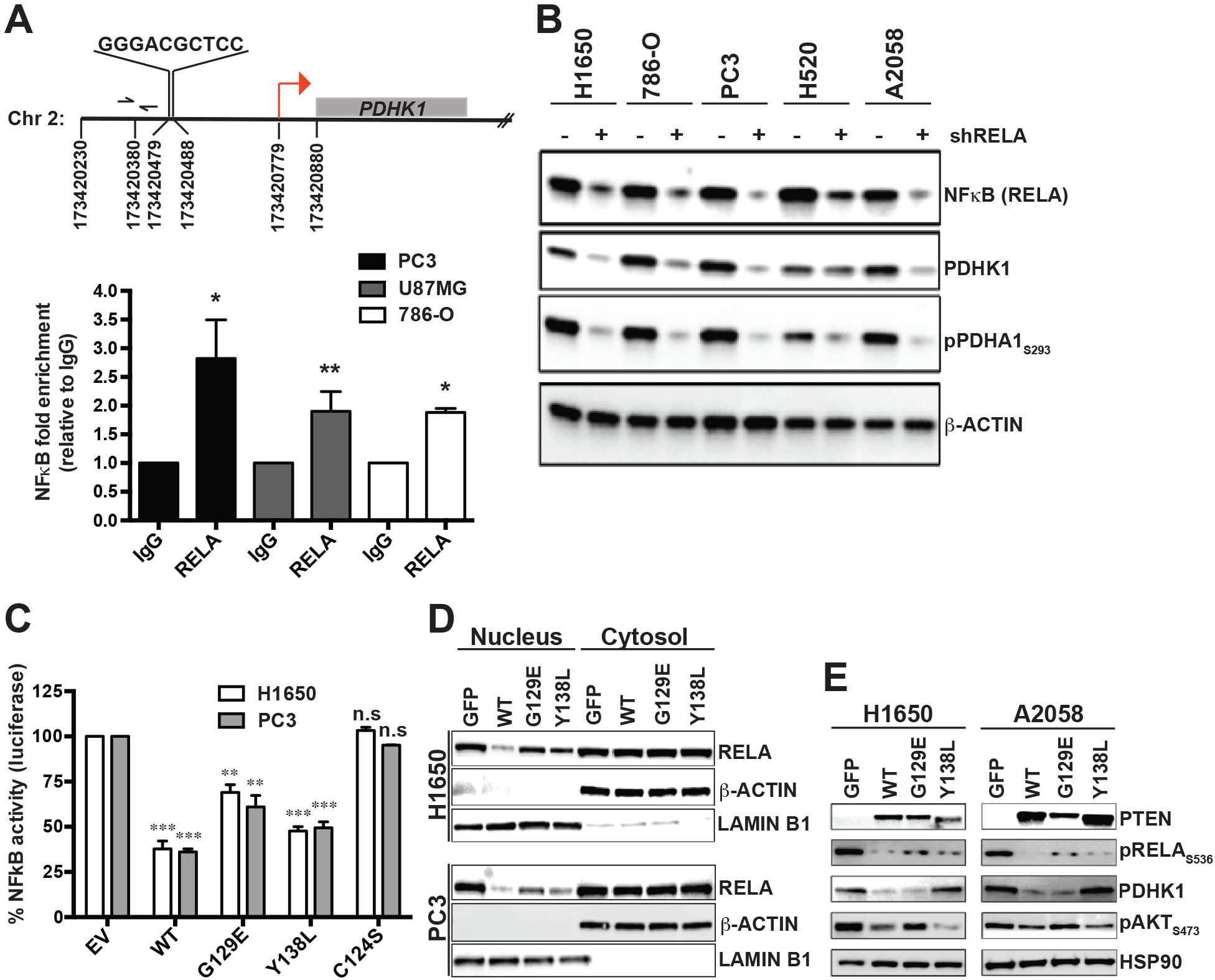
PTEN protein phosphatase and not lipid phosphatase downregulates PDHK1 by suppressing NFκB activation. (A) (Top) Identification of NFκB consensus binding site in the promoter of *PDHK1* gene located between nucleotide positions 173420479 and 173420488 in chromosome 2, ~300 bp upstream of the transcription start site (TSS, red arrow) at nucleotide position 173420779. (Bottom) NFκB (RELA) recruitment at PDHK1 promoter in PTEN-deficient cancer cells by ChIP assay. Primers (forward and reverse arrows) used to amplify a 118 bp region (spanning nucleotide position 173420380) ~40 bp upstream of the NFκB binding site in the *PDHK1* promoter are shown. Fold enrichment [RELA ChIP DNA (pg) to IgG DNA (pg)] data are shown as mean ± SEM (n = 3 replicates). \*\**p* < 0.01; \**p* < 0.05 compared to ‘IgG control’ by two-tailed unpaired t test with Welch’s correction. See also Table S2. (B) Western blots showing NFκB (RELA), PDHK1 and phospho-PDHA1 expression in PTEN-deficient cancer cell lines with or without stable *RELA* knockdown. shRELA, shRNA to RELA. (C) Effects of PTEN^WT^ or PTEN^G129E^ or PTEN^Y138L^ or empty vector (EV) expression in PTEN-deficient cancer cell lines on NFκB activity by luciferase reporter assays (Methods) are shown. Data are shown as mean ± SD (n = 2 replicates). \*\*\**p* < 0.001, \*\**p* < 0.01 compared to ‘empty vector control expressing PTEN-deficient cells’ by Tukey’s multiple comparisons one-way ANOVA test. (D) Effects of PTEN^WT^ or PTEN^G129E^ or PTEN^Y138L^ or GFP expression in PTEN-deficient cancer cell lines on NFκB (RELA) subcellular localization by nuclear-cytoplasmic fractionation and immunoblotting are shown. Western blots were also probed with anti-LaminB1 and anti-actinβ antibodies as nuclear and cytoplasmic markers, respectively. (E) Western blots showing PTEN, phospho-RELA, PDHK1 and phospho-AKT expression in PTEN-deficient stable cancer cell lines expressing PTEN^WT^, or PTEN^G129E^, or PTEN^138L^, or GFP. See also Figure S8 and table S3.

Consistent with this hypothesis, we found NFκB transcription factor RELA (p65) subunit recruitment at the PDHK1 promoter in multiple PTEN-deficient cancer cell types by chromatin immunoprecipitation (ChIP) and real time quantitative PCR analysis (Figure 4A). Further, we found NFκB suppression, using either an shRNA to knockdown *RELA* or the established NFκB small molecule inhibitor PBS-1086 (Blakely et al., 2015; Fabre et al., 2012) that acts as a specific inhibitor of RELA/B DNA binding and NFκB transcriptional activity inhibited NFκB, decreased PDHK1 expression, and suppressed phosphorylation of the PDHK1 protein substrate PDHA1 in a panel of PTEN-deficient cell lines (Figures 4B and S8C). Although prior studies have shown that PTEN loss, and consequent AKT activation can activate NFκB (Chiao and Ling, 2011; Dan et al., 2008; Gustin et al., 2001; Koul et al., 2001; Mayo et al., 2002), our data reveal that NFκB hyperactivation upon PTEN loss promotes PDHK1 expression and establish PDHK1 as a NFκB target gene.

Furthermore, NFκB inhibition by the established NFκB inhibitor PBS-1086 (Blakely et al., 2015; Fabre et al., 2012) also decreased cell survival specifically in PTEN-deficient cells (Figure S8D), phenocopying the effects of PDHK1 inhibition (Figure 2). These data suggest that PTEN loss renders cells dependent on PDHK1 for survival via NFκB activation.

We found that both the lipid, as expected (Gustin et al., 2001), and protein phosphatase activities of PTEN decreased NFκB activation, by using an established NFκB activation luciferase-reporter assay (Aoki and Kao, 1997; Blakely et al., 2015) and also by measurement of NFκB (RELA or p65) nuclear localization (Figures 4C and 4D). However, while the lipid phosphatase activity of PTEN specifically suppressed AKT activation (measured by phospho-AKT levels), as expected (Gustin et al., 2001), the protein phosphatase activity of PTEN was specifically required to suppress PDHK1 expression via decreased NFκB activation (measured by phospho-RELA levels) (Figure 4E). The collective data suggest that while both the lipid and protein phosphatase activities of PTEN regulate NFκB, the expression of PDHK1 is predominantly regulated by the protein phosphatase activity of PTEN via NFκB. These findings reveal the PTEN protein phosphatase activity as a regulator of both NFκB and PDHK1 and offer insights into PTEN’s activity-specific (lipid or protein phosphatase) modulation of cell signaling, growth, and metabolic phenotypes. These intriguing data show that differential transcriptional outputs of NFκB can be achieved by distinct upstream signaling events, an area for which there is some precedence and that will require future study (please see **Discussion**).

### PTEN protein phosphatase regulates NFκB and PDHK1 via NFκB activating protein (NKAP)

We next investigated how the PTEN protein phosphatase regulates its downstream effectors NFκB and PDHK1. We reasoned that de-phosphorylation of a specific protein target of the PTEN protein phosphatase activity could link PTEN to downstream NFκB and PDHK1 regulation. The full compendium of PTEN protein or lipid phosphatase specific effectors and substrates remains incompletely characterized. Therefore, to identify the potential PTEN protein phosphatase-specific effector that links PTEN to downstream NFκB and PDHK1 regulation, we used global (unbiased) mass spectrometry-based phospho-proteomics profiling (Beltrao et al., 2012; Jager et al., 2012) in a genetically-controlled system of PTEN-deficient cell lines with stable GFP or PTEN^WT^ or PTEN^G129E^ (lipid phosphatase mutant) or PTEN^Y138L^ (protein phosphatase mutant) re-expression (Figure 5A). We identified phospho-peptides and proteins that were significantly differentially regulated (Log2 fold-change < −0.5, *q* < 0.05) by each PTEN phosphatase activity, in comparison to the GFP, PTEN-null control (Figure 5A). As PDHK1 was regulated by the protein-phosphatase activity of PTEN across a range of PTEN-deficient models (Figures 3A, S7A and 4E), we prioritized the phospho-proteins that exhibited conserved and specific regulation by the PTEN protein-phosphatase activity in both H1650 (lung adenocarcinoma, *EGFR*-mutant) and A2058 (melanoma, *BRAF*-mutant) cells. The different tissue type and genetic background of these two cell lines provided an additional filter to identify a potentially conserved target. We found forty-two proteins exhibited PTEN protein phosphatase-specific regulation in both cell lines, including NF<u>κ</u>B activating protein (NKAP) (Figures 5A and S9A).

**Figure 5.**
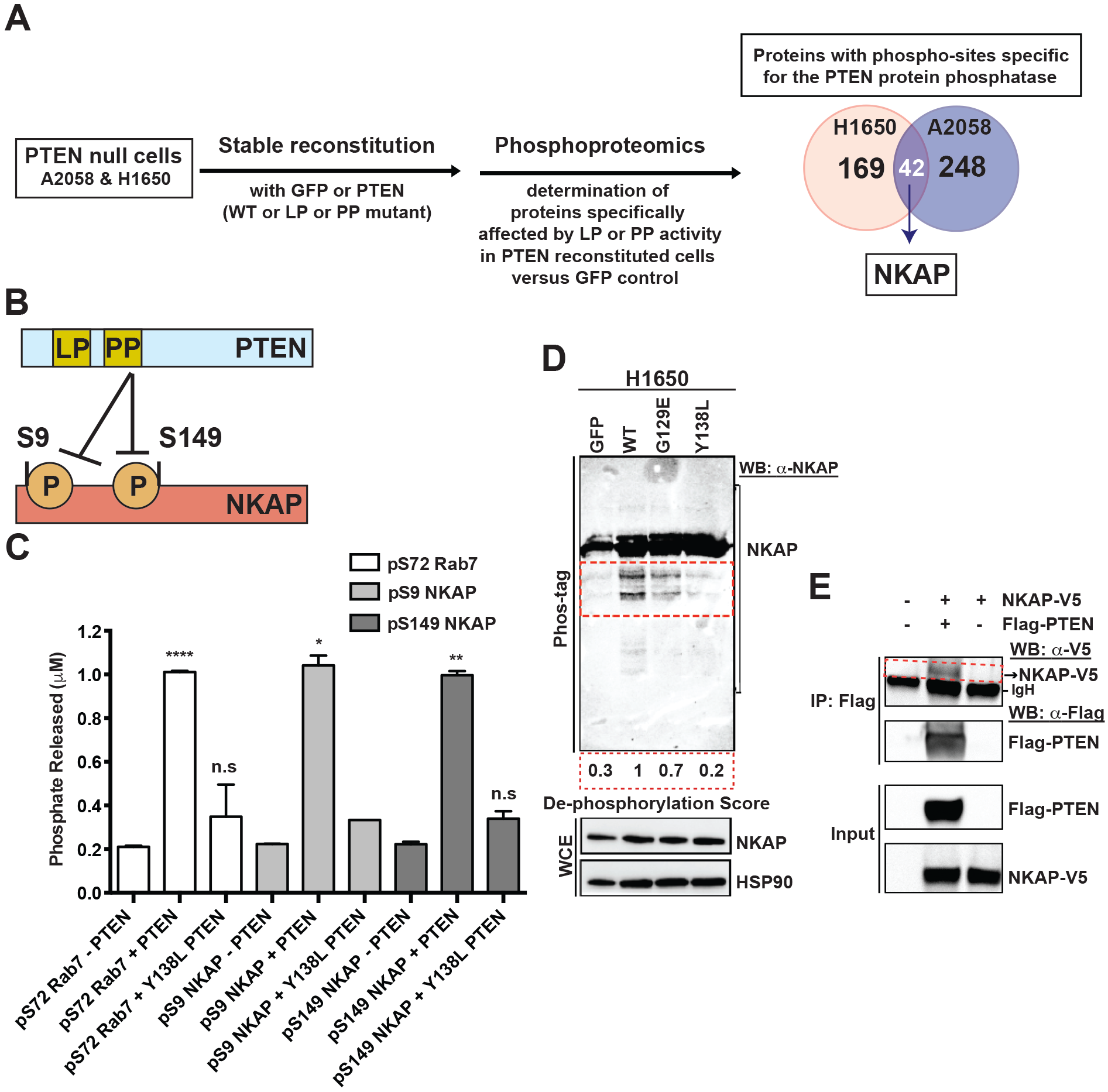
PTEN dephosphorylates NFκB activating protein, NKAP. (A) (Left) Workflow of the unbiased, global phospho-proteomic profiling used in PTEN-deficient cancer cell lines stably expressing PTEN^WT^ or PTEN^G129E^ or PTEN^Y138L^ or GFP to identify the phospho-peptides (and corresponding proteins) affected specifically by the protein or lipid phosphatase (PP or LP) activity of PTEN. (Right) Venn-diagram showing the number of proteins with phospho-sites specifically affected by the protein phosphatase activity of PTEN in H1650 (n = 169) and A2058 (n = 248) cells, with the phospho-proteins (including NFκB activating protein, NKAP highlighted in white) affected in both cell lines shown in the overlap (n = 42). (B) Schematic representation of NKAP phospho-sites at serines 9 and 149 affected by the protein phosphatase activity of PTEN. (C) Phosphate released (μM) from a phospho-S9-NKAP or phospho-S149-NKAP or phospho-S72-Rab7 (control) peptide after incubation without or with recombinant WT PTEN or mutant Y138L PTEN enzyme in an *in-vitro* Malachite green based colorimetric assay. Data are shown as mean ± SD (n=2 replicates). \**p* < 0.05; \*\**p* < 0.01; \*\*\*\**p* < 0.0001 compared to ‘no PTEN’ control by two-tailed unpaired t test with Welch’s correction. (D) Detection of phospho-NKAP and its de-phosphorylated species (indicated by the red dotted inset) in PTEN-deficient cancer cell lines stably expressing either PTEN^WT^ or PTEN^G129E^ or PTEN^Y138L^ or GFP by Phos-tag PAGE and immunoblotting. De-phosphorylation score indicates the extent to which expression of each PTEN mutant in PTEN-deficient cells suppresses NKAP de-phosphorylation relative to WT PTEN (set at 1). A lower de-phosphorylation score indicates less de-phosphorylation of NKAP. (E) Co-immunoprecipitation of NKAP (indicated by the red dotted inset) with PTEN upon over-expression of both NKAP-V5 and FLAG-PTEN and not NKAP-V5 over-expression alone or no over-expression in 293T cells followed by IP-FLAG is shown. Arrowhead denotes NKAP-V5; while the dark band below is background due to secondary antibody cross-reactivity to the immunoglobulin heavy chain (IgH) used in IP. See also Figure S9.

NKAP, which was originally identified as a regulator of NF<u>κ</u>B activation (Chen et al., 2003) (and hence the name), has various cellular functions including regulating Notch signaling and T-cell development (Pajerowski et al., 2009; Thapa et al., 2016). NKAP also interacts with RNA and RNA-binding proteins and plays a role in RNA splicing and processing (Burgute et al., 2014), chromosome segregation, and chromosome alignment through anchoring of CENP-E to kinetochores (Li et al., 2016). However, NKAP is not known to be linked to PTEN or the regulation of cellular metabolism or cancer. Given NKAP’s established ability to activate NFκB (Chen et al., 2003), which our data show directly regulates PDHK1 expression (Figure 4), NKAP was prioritized as a prime candidate PTEN effector arising from our phospho-proteomics screen (Figures 5A and S9A).

To investigate the hypothesis that NKAP links PTEN and PDHK1, first we examined whether NKAP is a phospho-protein target of PTEN in biochemical assays. The phospho-proteomics profiling revealed the predominant phosphorylation sites in NKAP in the two cell lines tested were at serines 9 and 149 (Figures 5B and S9A), consistent with other published (albeit previously un-confirmed) protein phospho-site mapping data (available at phosphosite.org). Using Ser 9 or Ser 149 phosphorylated NKAP peptides in an established Malachite-green based *in-vitro* phosphatase assay with recombinant PTEN^WT^ and PTEN^Y138L^ enzymes, we found that wild-type PTEN enzyme dephosphorylated NKAP at serines 9 and 149 with similar efficiency (as quantified by the amount of Pi released) as a known phospho-serine substrate of PTEN, phospho-S72 Rab7 (Shinde and Maddika, 2016), used as a positive control. The protein phosphatase defective PTEN (PTEN^Y138L^) enzyme had little or no activity for either phospho-NKAP or phospho-Rab7 and was comparable to the ‘no enzyme’ control (Figure 5C).

Further, by established phosphate affinity (Phos-tag) PAGE analysis (Kinoshita et al., 2009) (Figure 5D), wherein de-phosphorylated NKAP migrates faster than its phosphorylated form, we found enhanced NKAP dephosphorylation in PTEN deficient cells stably expressing PTEN^WT^ or PTEN^G129E^ (lipid phosphatase mutant) in comparison to those with stable PTEN^Y138L^ (protein phosphatase mutant) or GFP expression. These cellular Phos-tag assays independently confirmed that the protein phosphatase activity of PTEN predominantly regulates NKAP phosphorylation (Figure 5D). We noted no significant effect of PTEN re-expression on NKAP protein levels in PTEN-deficient cells (Figure 5D), suggesting that PTEN primarily regulates NKAP phosphorylation and not its expression. Additionally, no effect of PTEN expression on PDHK1 phosphorylation was observed (Figure S9A), suggesting that while PTEN regulates PDHK1 gene expression via NFκB (and consequently PDHK1 protein expression) it does not dephosphorylate PDHK1 protein.

Furthermore, NKAP co-immunoprecipitated with PTEN when over-expressed in 293T cells (Figure 5E, red box) under identical conditions in which the known PTEN-interactor NHERF2 (Takahashi et al., 2006) was co-immunoprecipitated with PTEN (Figure S9B). Consistent with these data, an independent affinity purification-mass spectrometry analysis (St-Denis et al., 2016) provided additional evidence for this PTEN-NKAP protein-protein interaction in cells. Collectively, these data reveal NKAP as a protein-interactor of PTEN and that PTEN dephosphorylates NKAP at two conserved serine residues S9 and S149.

To further investigate whether NKAP is a functionally important molecular link between PTEN and PDHK1, we examined whether NFκB activation and PDHK1 expression are each suppressed by silencing NKAP in PTEN-deficient cells. Indeed, we found that *NKAP* knockdown in different PTEN-deficient cell lines suppressed NFκB transcriptional activity using an established NFκB activation luciferase-reporter assay (Aoki and Kao, 1997; Blakely et al., 2015) (Figure 6A) and decreased NFκB activation (measured by p-RELA levels), without impacting pAKT levels (Figure 6C), consistent with prior work (Thapa et al., 2016). In concordance with our observation that NKAP silencing in the different PTEN-deficient cell lines suppressed NFκB activation, *NKAP* knockdown also suppressed transcriptional activation of the PDHK1 promoter (Figure 6B) and decreased expression of PDHK1 (Figure 6C), which we identified as a NFκB target gene (Figure 4). The collective findings indicate that loss of the PTEN protein phosphatase induces increased NKAP phosphorylation (on S9 and S149) and activates NFκB to upregulate PDHK1 expression, in a PI3K-AKT-independent manner.

**Figure 6.**
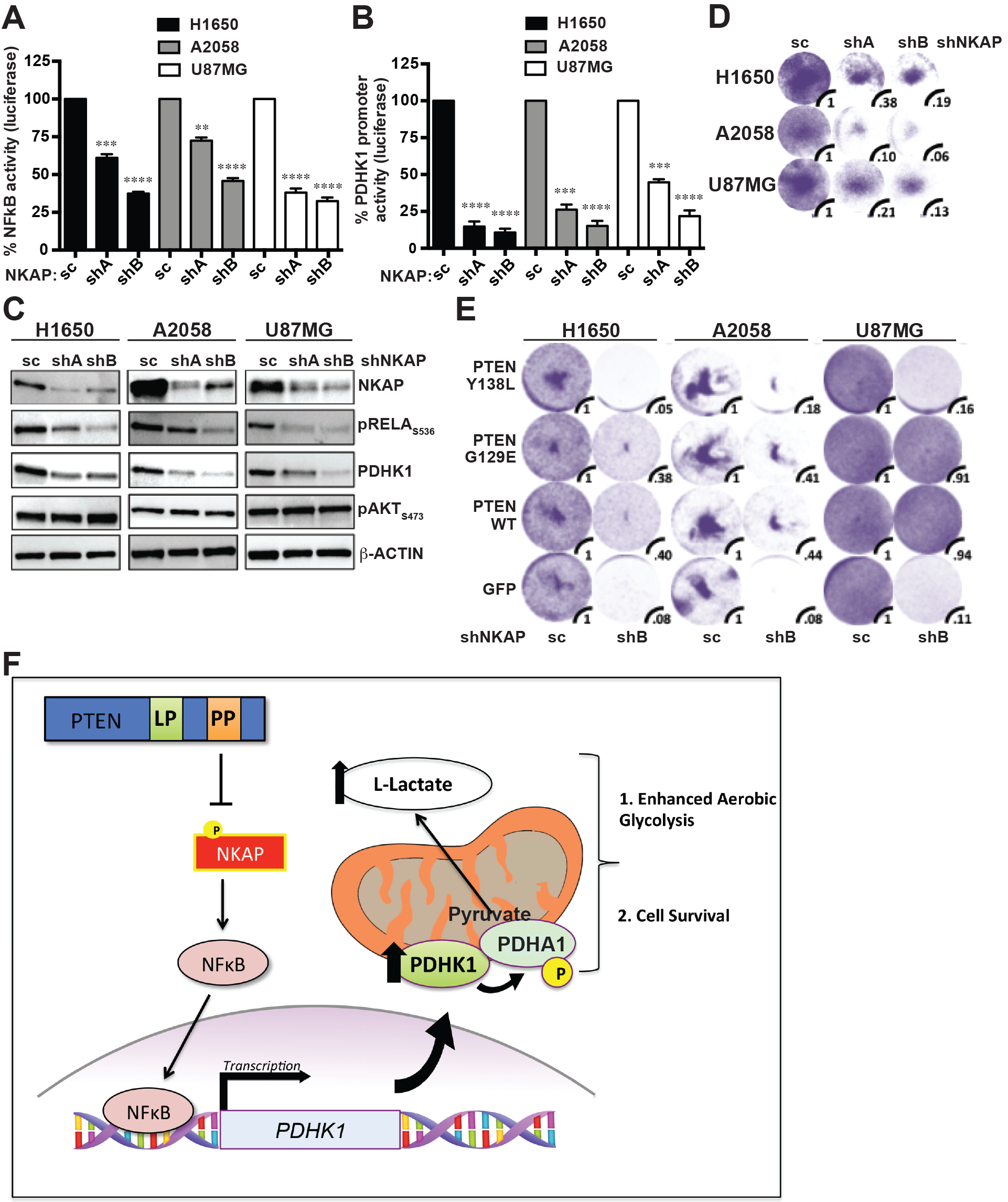
Depletion of NKAP decreases NFκB activation and PDHK1 expression and induces synthetic lethality specifically upon PTEN protein phosphatase loss. (A-B) Effects of stable *NKAP* knockdown in PTEN-deficient cancer cell lines on NFκB activity (A) or PDHK1 promoter activation (B) by luciferase reporter assays (Methods) are shown. shA and shB, shRNAs to NKAP and SC, scrambled control shRNA. Data are shown as mean ± SEM (n = 3 replicates). \*\*\*\**p* < 0.0001; \*\*\**p* < 0.001; \*\**p* < 0.01 compared to ‘scrambled control shRNA expressing PTEN-deficient cells’ by Tukey’s multiple comparisons one-way ANOVA test. (C) Western blots showing phospho-RELA, PDHK1 and phospho-AKT expression in PTEN-deficient cancer cell lines with or without stable *NKAP* knockdown. shA and shB, shRNAs to NKAP and SC, scrambled control shRNA. (D-E) Effects of stable *NKAP* knockdown in PTEN-deficient cancer cell lines without (D) or with stable PTEN^WT^ or PTEN^G129E^ or PTEN^Y138L^ or GFP expression (E) on cell growth by crystal violet staining assay are shown, with quantification of cell viability under each condition relative to cells expressing the scrambled control shRNA. shA and shB, shRNAs to NKAP and SC, scrambled control shRNA. (F) Model of cellular survival and energy metabolism regulation specifically by the protein-phosphatase activity of PTEN via a NKAP-NFκB-PDHK1-driven signaling axis. Loss of the PTEN protein-phosphatase activity promotes NKAP phosphorylation, NFκB activation, and PDHK1 upregulation, thereby enhancing aerobic glycolysis and rendering PTEN protein-phosphatase deficient cells dependent on NKAP and PDHK1 for survival.

Based on our findings, we propose a model in which specific loss of the PTEN protein phosphatase activates a downstream NKAP-NFκB signaling axis to drive PDHK1 expression and cellular dependence on PDHK1 for survival (Figure 6F). Consistent with this model, *NKAP* knockdown significantly decreased cell viability in PTEN-deficient, but not PTEN-expressing cells (Figure 6D), phenocopying the effects of both PDHK1 and NFκB inhibition in PTEN-deficient cells (Figures 2, S4 and S8C). Further, similar to PDHK1 inhibition (Figure 3D), NKAP silencing also significantly decreased the viability of cells specifically lacking the PTEN protein phosphatase activity (Figure 6E). PTEN-deficient cells re-constituted with PTEN^Y138L^ (protein phosphatase mutant) were generally as sensitive as PTEN-deficient cells to *NKAP* knockdown (Figure 6E). In contrast, cells re-constituted with PTEN that harbors the protein phosphatase activity (either PTEN^WT^ or PTEN^G129E^, the lipid phosphatase mutant) were generally less (or not at all) affected by NKAP suppression (Figure 6E). Together, these data suggest a critical role for the PTEN protein phosphatase in controlling cell survival via regulation of a PTEN/NKAP/NFκB/PDHK1 signaling axis and that interrupting this axis by suppressing either NKAP, NFκB, or PDHK1 in PTEN-deficient cells confers synthetic lethality.

## DISCUSSION

Our findings highlight an intricate link between cell signaling pathways and metabolic networks. Our data collectively provide insights into the diverse biological roles of the bi-functional PTEN phosphatase and unveil a synthetic essentiality of the metabolic kinase PDHK1 upon loss of PTEN in mammalian cells and in cancer.

PTEN is a tumor suppressor and a well-known negative regulator of the PI3K/AKT signaling pathway (Myers et al., 1998; Stambolic et al., 1998). While PTEN acts as a lipid phosphatase that dephosphorylates PIP3 to PIP2 at the cell membrane, PTEN was initially identified as a protein tyrosine phosphatase and later shown to be a dual-specificity (serine/threonine and tyrosine) protein phosphatase (Li et al., 1997; Maehama and Dixon, 1999). The biological role of the protein phosphatase activity of PTEN is less well studied than the lipid phosphatase activity of PTEN. Our study unveils functionally important molecular events that are specifically regulated by the protein phosphatase activity of PTEN in mammalian cells and cancer, thereby uncovering a potential therapeutic vulnerability in PTEN-deficient cancers.

We show that PDHK1 is a PI3K/AKT-independent effector of PTEN and establish a coordinated signaling pathway that operates specifically downstream of PTEN protein phosphatase to regulate PDHK1 in mammalian cells. Our study reveals an unanticipated biological role of the PTEN protein phosphatase in controlling cellular energy metabolism and cell survival via regulation of PDHK1. Furthermore, we identify a unique dependence of PTEN-deficient cells on PDHK1 for survival. Overall, our findings illustrate that specific loss of the protein phosphatase activity of PTEN results in NKAP-mediated NFκB activation, and consequent PDHK1 upregulation that promotes aerobic glycolysis and renders PTEN-deficient cells dependent on PDHK1 for survival (Figure 6F).

Our data suggest that PTEN’s different enzymatic activities can be used by cells to impart specificity in the regulation of cell signaling and transcriptional output (here, via NKAP/NFκB/PDHK1). Specifically, we confirm prior work (Dan et al., 2008; Gustin et al., 2001; Koul et al., 2001; Mayo et al., 2002) that loss of PTEN activates both PI3K/AKT and NFκB signaling. However, our data reveal that NFκB activation can occur upon loss of the PTEN protein phosphatase as well; indeed, increased PI3K/AKT signaling that occurs upon PTEN loss was not essential for driving downstream PDHK1 expression and cellular dependence. Our data suggest that certain factors such as the PTEN protein phosphatase substrate we identify here, NKAP, may impart such specificity in the context of PTEN protein phosphatase-mediated regulation of gene expression (e.g. PDHK1). The differential recruitment of NFκB to genomic loci such as the PDHK1 promoter may be guided not only by the underlying DNA sequence in canonical gene regulatory elements (e.g. promoter regions), but also by co-factor proteins (some with no known DNA binding domains) that influence preferential recruitment of NFκB to one target gene promoter (e.g. PDHK1 promoter) or another. In this way, the specific transcriptional competence of promoter and local chromatin regulatory elements may be conditioned by certain proteins that are not canonical transcription factors *perse*, such as NKAP. In support of this model, an increasing body of literature illustrates that different co-factor proteins (REL subunit-associating proteins, several of which have no DNA binding capability themselves) regulate NFκB binding at specific target genes under certain conditions (Wan and Lenardo, 2009). Consistent with this model, DNA binding specificities of other transcription factors at individual target gene promoters, such as the yeast Cbf1 (Siggers et al., 2011) and the drosophila HOX (Slattery et al., 2011) transcription factors, are also modulated by co-factor proteins (Met28 and EXD, respectively) that lack any DNA binding specificity or capacity themselves. Further, the interaction of a transcription factor with specific co-factors also depends on a specific stimulus. For example, the interaction of the estrogen receptor (ER) transcription factor with some of its cofactors is differentially modulated by various ligands (Ozers et al., 2005; Weatherman et al., 2002). Therefore, it is possible that NKAP, with no known conserved domains including DNA binding domains, is a co-factor that is specifically regulated by the PTEN protein phosphatase (and not lipid phosphatase) activity to direct differential NFκB promoter binding (e.g. to *PDHK1* and potentially other target gene promoters) only upon specific loss of the PTEN protein phosphatase. The proposed model may help explain how cells tightly control specificity in the differential expression of NFκB’s broad transcriptional repertoire by PTEN’s protein phosphatase activity or its lipid phosphatase activity in different cellular and biological contexts, an area for future investigation in both cancer and normal cells.

The significance of the cellular dependence on PDHK1 that we identify here is two-fold: it represents a previously unappreciated selective vulnerability in PTEN-deficient cells. Further, it exists in a wide range of PTEN-deficient cancer cell types as well as in normal cells with PTEN inactivation which reflects the generality of the conditional synthetic essentiality of PDHK1. Together, these findings raise the possibility that PDHK1 is a potential therapeutic target for evaluation in the large number of aggressive PTEN-deficient cancers for which there currently is no effective therapeutic strategy in the clinic, as inhibitors of PI3K signaling have not proven to be widely effective against PTEN-deficient cancers (Chandarlapaty et al., 2011; Ghosh et al., 2013; Rodon et al., 2013). Moreover, additional clinical relevance of our findings is supported by our data showing that PDHK1 expression levels were increased in PTEN-deficient human tumors and high PDHK1 expression was a biomarker of inferior patient survival in this class of patients. Thus, increased PDHK1 could be used to help identify PTEN-deficient cancer patients that may benefit from tailored and more aggressive cancer surveillance and/or treatment.

Broadly, our findings illustrate the importance of defining activity-specific PTEN functions that contribute to physiologic regulation in normal cells and that when dysregulated contribute to genetically-driven human diseases such as cancer.

## Acknowledgements

The authors would like to thank all the members of the Bivona laboratory for critical review of the manuscript and acknowledge funding support from the following sources to (T.G.B): NIH Director’s New Innovator Award, Howard Hughes Medical Institute, Doris Duke Charitable Foundation, American Lung Association, National Lung Cancer Partnership, Sidney Kimmel Foundation for Cancer Research, Searle Scholars Program, California Institute for Quantitative Biosciences, QB3 (to T.G.B and N.J.K) and Ruth L. Kirschstein National Research Service T32 Award in molecular & cellular mechanisms in cancer (to N.C). We thank Timothy Fouts and Jeffrey Meshulam at Profectus Biosciences, Inc. for generously providing PBS-1086. We thank Bhagyashri Burgute and Angelika Noegel at University of Cologne, Germany for generously providing the NKAP mouse mAb K85-80.

## Author Contributions

N.C. and E.P. designed and performed experiments and analyzed data. N.C. and B.B. conducted PTEN purification. G.H., L.L., and C.H.E., J.F., E.C., M.K.M. performed experiments and analyzed results. V.O. performed animal and IHC studies. J.J.Y. performed ChIP experiments and sequencing analysis. S.A. analyzed molecular profiling data. J.S. performed single-cell and high throughput fluorescence imaging and analyzed data. T.I. and M.H. performed analysis and interpretation of TCGA tumor data. E.V., J.R.J., B.W.N., J.V.D., N.J.K. conducted and analyzed proteomics experiments. N.C., E.P., A.J.S, T.G.B. wrote the manuscript with input from all authors.

## METHOD DETAILS

### Cell Lines, antibodies and reagents

Human lung adenocarcinoma cell lines (H1650, HCC827, H1975) were generously provided by Dr. William Pao (Vanderbilt University). PC3 cells were a kind gift from Dr. Stefan Grimm (Imperial College London), U87MG cells were kindly provided by Dr. William A. Weiss (UCSF) and 786-O cells from Dr. Celeste Simon (University of Pennsylvania). All other cells lines were obtained from American Type Culture Collection (ATCC).

RelA, p-RelA^S536^, HK2, PTEN, AKT, pAKT^S473^, GAPDH, HSP90, H3, cleaved-PARP, p-S6^S240^, p-GSK-3α/β^S21/9^ and GST antibodies were obtained from Cell Signaling Technologies, PDHK1 antibody was from Enzo Life sciences, p-PDHA1^S293^ antibody was from Millipore, FLAG and CA9 antibodies were from SIGMA, V5 antibody was from Thermo Fisher and NKAP polyclonal, HIF1, PDHK2, PDHK3 and PDHK4 antibodies were from Abcam. HRP-conjugated anti-Mouse and anti-Rabbit secondary antibodies were obtained from Biorad and Cell Signaling Technologies.

Phospho-peptides pS9-NKAP (SGSRpSPDREA), pS149-NKAP (VWGLpSPKNPE) and p-S72-Rab7 (ERFQpSLGVAF) were purchased from Elim Biopharmaceuticals, Hayward, CA.

Drugs were re-suspended in DMSO at 10 mM final concentration and stored at −20 °C. DCA dilutions were made fresh just before use in water.

### Plasmids, stable cell lines and transfection

Mammalian retroviral expression plasmids pBABE-PTEN^WT^ (Cat# 10785), pBABE-PTEN^C124S^ (Cat# 10931), pBABE-PTEN^G129E^ (Cat# 10771), pBABE-Puro (Cat# 1764) and pBABE-GFP (Cat# 10668) were purchased from Addgene (Cambridge, MA). pBABE-PTEN^Y138L^ expression plasmid was engineered using standard molecular biology techniques. Mammalian expression plasmid pCMV-2xFLAG-PTEN^wt^ (Cat# 28298) was purchased from Addgene (Cambridge, MA) and mammalian lentiviral expression plasmid pLX304-NKAP-V5 was purchased from Dharmacon (Cat# OHS6085). pGreenFire1-NF-kB (EF1α-Puro) lentiviral plasmid (Cat# TR012PA-P) coding for the NF-κB luciferase reporter was purchased from SBI (System Biosciences). *PDK1* promoter reporter construct (Cat# S721832) coding for the PDHK1 promoter luciferase reporter was purchased from SwitchGear Genomics.

293GPG retroviral packaging cells(Ory et al., 1996) were transfected with pBABE (empty vector) and pBABE-PTEN^WT^ or -PTEN^Mutant^ constructs using Lipofectamine-2000 (Invitrogen) according to manufacturer’s instructions. Virus containing media was harvested three days post transfection and used to infect the indicated cell lines. Cells were incubated with virus containing media supplemented with 6 μg/ml of polybrene for 24 hours. Media was changed to standard cell growth media (RPMI-1640 or DMEM + 10% fetal bovine serum and 100 U/ml penicillin G and 100 U/ml streptomycin sulphate) and cells were expanded for 48 hours, at which point puromycin (0.5-2 pμg/ml) was added to the media and cells were allowed to grow for an additional 4 days. Cells that survived puromycin selection (stable cell lines) were used in all subsequent experiments.

High titer pre-made adenovirus (Cat# 000028A) expressing PTEN was purchased from ABM (Applied Biological Materials), Richmond, Canada and used for transduction of PTEN-deficient cells for transient PTEN over-expression following manufacturer’s instructions.

HEK 293T cells were transfected with pCMV-2xFLAG PTEN^WT^ and pLX-TRC304-NKAP-V5 expression plasmids using TransIT-LT1 transfection reagent (Mirus Bio) according to manufacturer’s instructions. Briefly, the plasmid diluted in serum-free OptiMEM medium was mixed with TranslT-LTl transfection reagent in 1:3 ratio. After incubating the DNA-TranslT-LT1 mixture at room temperature for 15 min, the complexes were added to cells to allow the transfection of plasmid.

### Gene expression profiling

Three biological replicate retroviral infections of H1650 cells were performed with pBABE-PTEN or pBABE-GFP as described above, followed by selection in 2 μg/ml puromycin for 2 days. Stable PTEN expression was confirmed by immunoblot analysis 5 days postinfection. Total RNA was isolated for each replicate using the RNeasy kit (Qiagen) according to manufacturer’s instructions. After quality control validation, 2 μg of total RNA was processed for hybridization and image quantification of U133A2.0 gene expression arrays (Affymetrix).

### Real time qRT-PCR

*PDHK1* gene expression changes observed by microarray analysis were validated by real-time qRT-PCR in paired cell lines engineered to stably lack or express PTEN^WT^. *PDHK1* gene expression regulation by different phosphatase activities of PTEN was also measured by realtime qRT-PCR in PTEN-deficient cell lines stably expressing PTEN^WT^, PTEN^G129E^, PTEN^Y138L^ or GFP. Briefly, for real time qRT-PCR analysis total RNA was extracted from cells using the RNeasy kit (QIAGEN) and cDNA synthesis was performed according to manufacturer’s instructions. Real time qRT-PCR was performed on the Quant Studio TM 12K Flex real-time qPCR system using Taqman probes (Applied Biosystems, Life Technologies) specific to the coding region of the *PDHK1* gene. GAPDH expression levels were used as an internal control for normalization of cDNA content. The QuantStudio 12K Flex V1.1.2 software was used to analyze real time qRT-PCR data.

### Immunohistochemistry

For immunohistochemistry, 5-15 μm thick formalin-fixed paraffin embedded (FFPE) human tissue sections were purchased from US Biomax (GBM: Cat# GL1921, Prostate adenocarcinoma: Cat# PR803a and Lung Squamous Cat# LC801) and stained with PTEN or PDHK1 antibodies following manufacturer’s instructions. Stained slides were digitized using the Aperio ScanScope slide scanner (Aperio Technologies) with a 40x objective. Scanned images from the immunohistochemistry staining were used to score expression of PTEN or PDHK1 in paired samples (scoring expression on a scale of 1 to 3, with 1 indicating ‘low’, 2 ‘intermediate’, and 3 ‘high’ expression of each protein). Bars indicate 50 μm and arrows indicate representative tumor cells scored for PTEN and PDHK1 expression.

### Cell viability and growth assays

Cells were seeded overnight at a density of 5,000 cells per well in 96-well plates in RPMI or DMEM containing 10% FBS and treated with indicated drugs for 72 hours. Viable cell numbers were determined using the CellTiter-Glo luminescent cell viability assay according to manufacturer’s instructions (Promega). Each assay consisted of eight replicate wells and was repeated at least in three independent experiments. Data are expressed as percentage of the cell viability of control cells. when Cell viability was measured in an ATP independent assay, cells were plated in 6-well plates and fixed with 4% paraformaldehyde. Crystal Violet was used to stain the viable cells and imaged using an ImageQuant LAS 4000 instrument (GE Healthcare). To quantify the number of viable cells, stained cells were solubilized in 1% SDS and absorbance was measured at 562 nm in a plate reader (SpectraMax M5, Molecular devices).

For the determination of rescue score, cells were treated with a dose-response of DCA and stained with crystal violet 72 hours after treatment. In each cell line, the degree of cell viability in the highest concentration of DCA (20 mM) was normalized to the vehicle control for each genetic background (cells expressing GFP or PTEN^WT^ or PTEN^Mutant^). This normalized cell viability value was then used to calculate the rescue score as: (normalized viability in the PTEN-reconstitution condition/ normalized viability in the PTEN-deficient, GFP, condition). A rescue score >1 indicates the degree of rescue from lethality upon PDHK1 inhibition.

For western blotting, cells were scraped and lysed in lysis buffer (50 mM Tris HCl pH 8.0, 150 mM sodium chloride, 0.1% SDS, 0.5% sodium deoxycholate, 1% Triton X 100, 5mM EDTA containing protease and phosphatase inhibitors (Roche Diagnostics) and clarified by sonication and centrifugation. After quantification of total protein by Pierce BCA assays (Thermo Scientific, Rockford, IL), equal amounts of total protein (10 μg) was separated by 415% SDS-PAGE, transferred to nitrocellulose membrane for further analysis by standard immunoblotting protocol using specific antibodies.

### L-lactate measurement

Extracellular L-lactate was measured using a lactate assay kit from Cayman Chemical Company (Cat# 700510) following manufacturer’s instructions. Briefly, cells were plated at equal numbers in at least 8 micro wells per condition and cultured with the same volume of media. The assay employs the feature of NAD^+^ reduction to NADH which occurs concomitantly with the oxidation of lactate to pyruvate. NADH reacts with the fluorescent substrate to produce high fluorescence at em540/ ex590 wavelength. The greater the signal the higher is the lactate concentration in the culture media. Each experiment was repeated in triplicate. For glucose conditioned media, RPMI or DMEM without glucose was used (Gibco) and 2DG (2-deoxyglucose) was supplemented at 10 mM final concentration.

### Colony formation assays

50,000 cells were seeded in 0.35% Noble-agar overlaying 0.6% Noble agar base in 60 mm dishes (Falcon). Cells were grown for 21-28 days and media was changed after every 4 days. Colonies were visualized after staining with 0.005% crystal violet, imaged and quantified.

### Animal studies

A2058-GFP and A2058-PTEN tumor xenografts were generated by injection of 2×10^6^ cells/ tumor mixed with matrigel in Scid mice. Stable PTEN expression in A2058 cells was confirmed by immunoblotting before *in-vivo* transplantation. Tumors were allowed to grow until they reached a minimum volume of 200 mm^3^ at Day 17 when both group of animals (with GFP and PTEN expressing tumors) received treatment with DCA (750 mg/ L) in drinking water. Tumor growth was assessed at the indicated time-points by caliper measurements over 31 days post *in-vivo* transplantation and data shown represent the endpoint of the study. A minimum of 6 tumors per treatment group was assessed for the duration of the study.

### RNA interference assays

pLKO.1-puro based shRNAs for *PDHK1*, *RelA* and *PTEN* were purchased from SIGMA. At least 2 separate constructs of pre-validated hairpins were used for each gene to exclude off-target effects. The shRNAs purchased from the TRC collection are the following: *PDHK1*: [TRCN0000194672, TRCN0000196891, TRCN0000196728, TRCN0000006261 (sh#1), TRCN0000006263 (sh#2)], *PTEN*: [TRCN0000002749, TRCN0000002748, TRCN0000002747, TRCN0000230370], *RELA*: [TRCN0000014683, TRCN0000014684, TRCN0000014686], *NKAP*: [TRCN0000145605, TRCN0000145475, TRCN0000144065, TRCN0000143845, TRCN0000143380]. sh#1, sh#2: shRNAs targeting the 3’UTR of *PDHK1*. Viral supernatant was produced following standard lentiviral production protocols using calcium phosphate transfection, 20 μg of pLKO.1-shRNA and 20 μg of ViraPower mixed packaging plasmids from Life technologies. Virulent media was used to infect the cells for 6 hours. Selection with puromycin was started 72 hours post infection, immunoblotting to confirm stable knockdown was done 5 days post infection and crystal violet staining to assess the effects of stable knockdown on cell viability was done 6 days post infection. Crystal violet staining to assess the effects of stable *PDHK1* knockdown in PTEN-deficient and -proficient cell lines on cell viability was done 9 days post infection. 96 hours post puromycin selection of cells for stable *PDHK1* knockdown, cells were transduced with adenovirus (ABM, Cat# 000581A) expressing shRNA-resistant *PDHK1* at MOI 1000. Crystal violet staining to assess the effects of stable *NKAP* knockdown in PTEN-deficient cell lines or in PTEN-deficient cell lines stably expressing either PTEN^WT^ or PTEN^G129E^ or PTEN^Y138L^ or GFP on cell viability was done 6 days post infection.

### Chromatin immunoprecipitation (ChIP) assays

ChIP assays were performed using the chromatin IP kit from Cell Signalling Technology (Cat#9003) following manufacturer’s instructions. Briefly, 4×10^7^ cells were treated with formaldehyde to crosslink proteins to DNA and nuclear/cytosolic fractionation was performed. Nuclear lysates were sonicated using the Covaris S2 sonicator and an aliquot of DNA was purified to check for sonication efficiency (fragments of ~500bp). Anti-p65, anti-IgG and anti-H3 (positive control) antibodies were used to pull down the indicated protein-bound chromatin. washing, elution and reverse crosslink followed by DNA purification concluded the sample preparation. Quantification of NFκB (or p65) enrichment was done using real-time qPCR with primers designed to amplify a 118 bp region spanning nucleotide position 173420380 in the *PDHK1* promoter (EpiTect ChIP qPCR Primer Assay for human *PDK1*, NM002610.3 (-)01Kb, Cat# GPH1007880(-)01A, QIAGEN. Assay position: Chromosome 2, 173420380, TSS position: Chromosome 2, 173420779, Assay tile: (-) 01Kb). A standard curve was generated to examine the efficiency of amplification by plotting Ct (threshold cycle) versus DNA quantity (Log10 scale). The fold enrichment was calculated using the following two formulas: Formula 1: Y = M(Log10X) + B (X=DNA quantity, Y=Ct, M=slope of the curve, B=Ct value where X=1). Formula 2: Fold enrichment = (ChIP DNA Quantity)/ (IgG DNA Quantity).

### Luciferase reporter assays

pGreenFire1-NF-kB (EF1α-Puro) lentiviral plasmid containing four NFκB transcription factor response elements fused in tandem upstream of a luciferase gene under a minimal CMV promoter was transfected into PTEN-deficient cancer cells stably expressing either PTEN^WT^ or PTEN^G129E^ or PTEN^Y138L^ or GFP to measure the effects of PTEN’s protein and / or lipid phosphatase activities on the transcriptional activity of NFκB. pGreenFire1-NF-kB (EF1α-Puro) lentiviral plasmid containing four NFκB transcription factor response elements was also transfected into PTEN-deficient cancer cell lines with stable *NKAP* knockdown to study the effects of NKAP downregulation on NFκB activity. *PDK1* promoter reporter plasmid containing the PDHK1 promoter fused to luciferase was transfected into PTEN-deficient cancer cell lines with stable *NKAP* knockdown to measure the effects of NKAP downregulation on PDHK1 promoter activation. Luciferase activity was measured 48 hours after transfection. Luciferase detection reagent was added (Bright-Glo, Promega) at 100 μL per well and luminescence was measured after 5 min at RT in a plate reader (SpectraMax M5, Molecular devices).

### Phosphopeptide screen set up and sample processing for mass spectrometry

GFP or PTEN-WT or PTEN-G129E or PTEN-Y138L was stably introduced in two different PTEN-deficient cancer cell lines H1650 and A2058 with different genetic and histologic backgrounds. Cells were lysed in a buffer containing 8M urea, 100 mM Tris.Cl pH 8.0, 150 mM NaCl, and protease inhibitors (EDTA free cOmplete cocktail tablet, Roche). Lysates were reduced with 4 mM TCEP for 30 minutes at room temperature and alkylated with 10 mM iodoacetamide for 30 minutes at room temperature in the dark. Samples were diluted 1:4 in 100 mM Tris pH 8.0 to reduce urea concentration to 2M and digested with trypsin (1:100 enzyme: substrate ratio) overnight at 37°C. Peptides were desalted with Sep-Pak reversed phase C18 solid phase extraction (Waters) according the manufacturer’s instructions and phosphopeptides were purified using an immobilized metal affinity chromatography approach. Purified phosphopeptides were analyzed in an LTQ Orbitrap Elite mass spectrometry system (Thermo Scientific) equipped with a Proxeon Easy nLC 1000 ultra high-pressure liquid chromatography and auto sampler system. Samples were injected onto a C18 column (25 cm × 75 um I.D. packed with ReproSil Pur C18 AQ 1.9 um particles) in 0.1% formic acid and then subjected to a 4-hour gradient from 0.1% formic acid to 30% ACN/0.1% formic acid. The mass spectrometer collected data in a data-dependent fashion, collecting one full scan in the Orbitrap at 120,000x resolution followed by 20 collision-induced dissociation MS/MS scans in the dual linear ion trap for the 20 most intense peaks from the full scan. Dynamic exclusion was enabled for 30 seconds with a repeat count of 1. Charge state screening was employed to reject analysis of singly charged species or species for which a charge could not be assigned.

### *In vitro* phosphatase assays

5-14 amino acid (SGSRpSPDREA) pS9-NKAP or 145-154 amino acid (VWGLpSPKNPE) pS149-NKAP or 68-77 amino acid (ERFQpSLGVAF) p-S72-Rab7 peptide (used as a +ve control) was incubated without or with bacterially expressed and purified (as described before(Taylor and Dixon, 2003)) WT PTEN or mutant Y138L PTEN enzyme in reaction buffer (25mM HEPES, pH 7.5, 10 mM MgCl2, 10 mM DTT) at 37°C for 90 min. Following incubation, the released phosphate from each phospho-peptide ± WT PTEN or mutant Y138L PTEN enzyme was detected using Malachite Green Assay Kit (#K1500, Echelon) by measuring the absorbance at 620 nm.

### Phos-tag assays

Phos-tag PAGE analysis was carried out to detect phosphorylated-NKAP and its de-phosphorylated species according to manufacturer’s instructions (Wako Chemicals, Japan). Briefly, PTEN-deficient cancer cells stably expressing either PTEN^WT^ or PTEN^G129E^ or PTEN^Y138L^ or GFP were lysed and subjected to Zinc-based polyacrylamide gel electrophoresis. After electrophoresis, Phos-tag acrylamide gels were washed with gentle shaking in transfer buffer containing 10 mM EDTA for 30 min and then incubated in transfer buffer without EDTA for 15 min. Proteins were then transferred to nitrocellulose membrane for further analysis by standard immunoblotting protocol using specific antibodies.

### Nuclear and cytoplasmic fractionation

Subcellular localization of NFκB (or p65) in PTEN-deficient cancer cells was performed as described before(Suzuki et al., 2010).

### Immunoprecipitation

For immunoprecipitation assays, 48 hours post-transfection of different plasmids in HEK 293T cells, the cells were resuspended in lysis buffer (200 mM NaCl, 0.5 mM EDTA, 0.5% NP40, 50 mM Tris.Cl, pH 7.5) containing 250 U Benzonase (SIGMA)/ 5 ml and protease inhibitors and lysed by a rapid freeze-thaw cycle (10 min on dry ice followed by 1 min at 37°C). The wholecell extracts (WCE) obtained by centrifugation were incubated overnight with M2 agarose-FLAG beads (SIGMA) at 4°C. The immunocomplexes were washed once with wash buffer I (50 mM Tris (pH 7.4, 150 mM NaCl, 0.05% NP40), three more times with wash buffer II (50 mM Tris (pH 7.4, 150 mM NaCl), eluted from the beads with elution buffer (150 ng/ μl FLAG peptide in wash buffer II) and applied to SDS-PAGE.

## QUANTIFICATION AND STATISTICAL ANALYSIS

### Statistical Analyses

Statistical differences between two experimental groups were calculated using the unpaired twotailed Student’s *t*-test. Statistical differences between multiple experimental groups were calculated using the multiple comparisons one-way ANOVA test. Statistical differences of tumor volumes were calculated using the two-way ANOVA test. A significance level of *p* < 0.05 or less was used throughout the study.

### TCGA Data Analysis

Data preparation: TCGA pan-cancer gene expression by RNAseq using the Illumina HiSeq technology was downloaded (https://toil.xenahubs.net/download/tcga_RSEM_Hugo_norm_count.gz). Data from all TCGA cohorts (N=10535) are combined to produce this dataset. Values are Log2 (x+1) transformed RSEM values.

TCGA pan-cancer gene-level copy number variation (CNV) estimated using the GISTIC2 threshold method was downloaded. Data from all TCGA cohorts (N = 10845) to produce this dataset. GISTIC2 further thresholded the estimated values to −2, −1, 0, 1, 2, representing homozygous deletion, single copy deletion, diploid normal copy, low-level copy number amplification, or high-level copy number amplification. Genes are mapped onto the human genome coordinates using UCSC cgData HUGO probeMap (https://tcga.xenahubs.net/download/TCGA.PANCAN.sampleMap/Gistic2_CopyNumber_Gistic2_all_thresholded.by_genes.gz).

Samples were subtyped based on *PTEN* copy number status, samples with CNV = 0 were assigned to *PTEN* normal (N = 6640), samples with CNV = −1 or −2 were assigned to *PTEN* copy number loss (N = 3422). *PDHK1* gene expression was extracted from TCGA pan-cancer normalized RNAseq RFKM matrix and matched with samples with *PTEN* normal group (N = 5995) or *PTEN* copy number loss group (N = 2845).

Expression analysis of patient samples: Expression levels of *PDHK1* mRNA after normalization were compared between patient samples with single or double copy number loss of *PTEN* to patients with copy number neutral *PTEN* by boxplot analysis. The minimum, first quartile, median, third quartile and maximum were shown with horizontal lines from lower to upper with 1.5 interquartile range. Unfilled circles are outlier expression of *PDHK1* in these two subgroups. Reported *p*-values are for two-tailed unpaired t test with Welch’s correction comparing the expression samples with *PTEN* copy number loss to the normal samples.

Survival analysis: Patient survival outcomes were downloaded from the Firehose web portal. We excluded from survival analysis patients over 75 years of age (N = 416) or patients who died less than 30 days from diagnosis (N = 184). Reported *p*-values are for a cox proportional hazards likelihood ratio test comparing a baseline model to a three-category model indicating a split of the patient cohort into three bins according to lower quartile (25 percentile and below), inter-quartile range (25-75 percentiles) and upper quartile (75 percentile and above) *PDHK1* normalized expression values.

### Gene Expression Array (Affymetrix) Data Analysis

Raw data was processed using R/Bioconductor package with Robust Multichip Average (RMA) normalization to identify differentially expressed genes between stable PTEN or GFP expressing H1650 cells. The normalized expression intensities were Log2 transformed and multiple t-test for null hypothesis was applied adjusted with Benjamini-Hochberg’s false discovery rate (FDR). The significantly differentially expressed genes were selected by applying fold change cut-off > 2, \**p* < 0.05 and FDR adjustedp-value or *q*-value < 0.2.

### Phosphopeptide Quantification and Data Analysis

Raw mass spectrometry data were analyzed using the MaxQuant software package (version 5). Data were matched to SwissProt reviewed entries for Homo sapiens in the UniProt protein database. MaxQuant was configured to generate and search against a reverse sequence database for false discovery rate calculations. Variable modifications were allowed for methionine oxidation, protein N-terminus acetylation, and serine, threonine, and tyrosine phosphorylation. A fixed modification was indicated for cysteine carbamidomethylation. Full trypsin specificity was required. The first search was performed with a mass accuracy of +/− 20 parts per million and the main search was performed with a mass accuracy of +/− 6 parts per million. A maximum of 5 modifications were allowed per peptide. A maximum of 2 missed cleavages were allowed. The maximum charge allowed was 7+. Individual peptide mass tolerances were allowed. For MS/MS matching, a mass tolerance of 0.5 Da was allowed and the top 6 peaks per 100 Da were analyzed. MS/MS matching was allowed for higher charge states, water and ammonia loss events. The data were filtered to obtain a peptide, protein, and site-level false discovery rate of 0.01. The minimum peptide length was 7 amino acids. Results were matched between runs with a time window of 2 minutes for technical duplicates. The label-free PTEN phosphoproteomics data was analyzed using an in-house computational pipeline built for the analysis of post-translationally modified peptides with mixed effect models, implemented in the MSstats (v2.3.4) Bioconductor package(Choi et al., 2014). First, protein identifiers were converted into modification site identifiers, contaminant and false positive MaxQuant search results were removed and all samples were normalized per cell line by median-centering the log2-transformed MS1-intensity distributions. Next, the MSstats group Comparison function was run with the following options: no interaction terms for missing values, no interference, unequal intensity feature variance, restricted technical and biological scope of replication. Statistically significant changing sites between PTEN variants and the GFP control were selected by applying a Log2 fold-change < − 0.5 and q-value < 0.05 threshold.

---

**Figure S1.**
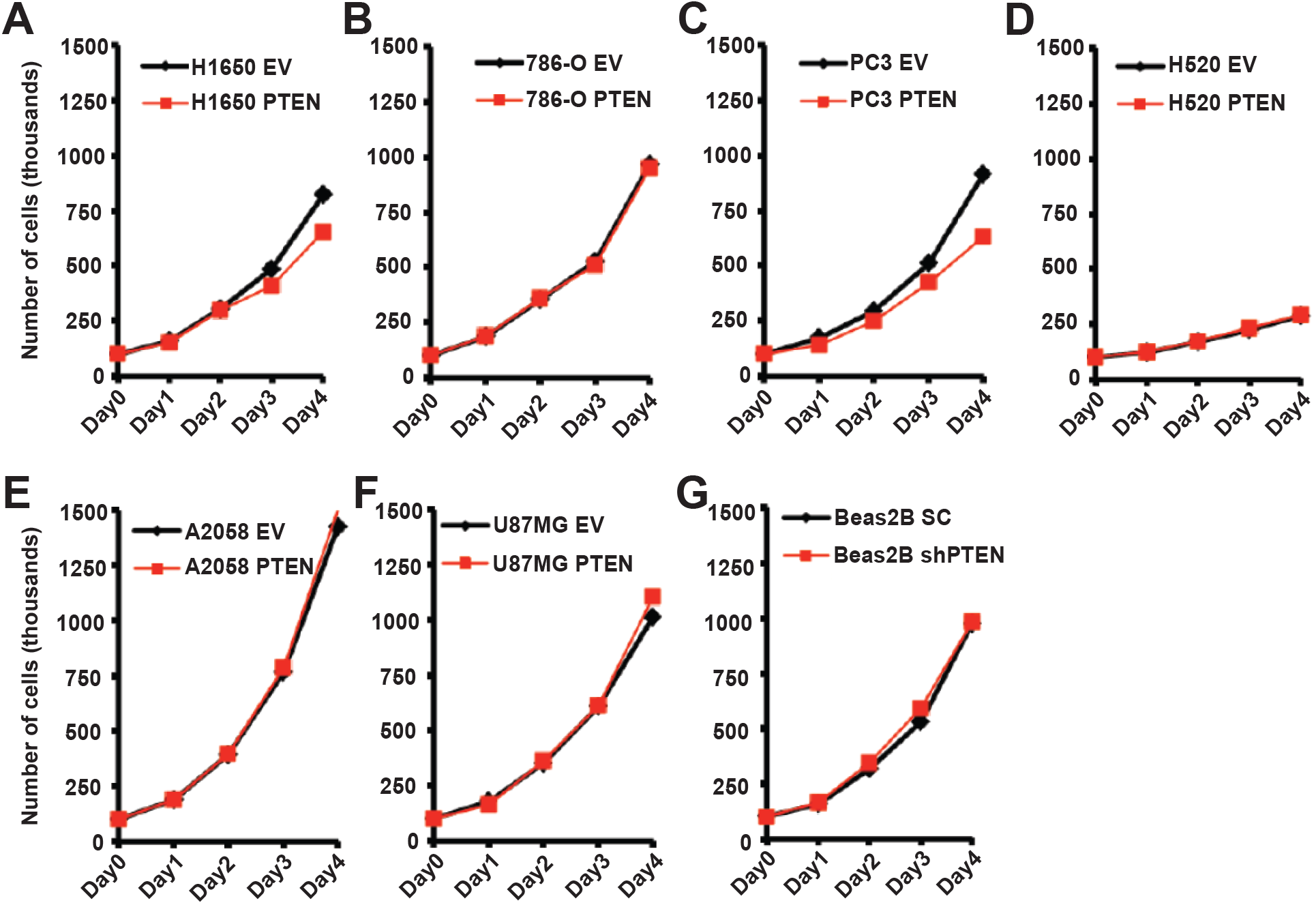
Related to Figure 1. PTEN-deficient cells with stable re-expression of PTEN or PTEN-proficient cells with stable *PTEN* knockdown grow similarly as parental cells. (A-F) The effects of stable PTEN expression or (G) stable *PTEN* knockdown in the indicated cancer and normal cell lines on cell growth are shown. Cells were plated at 100,000 cells per condition at Day 0, cell number was enumerated at each time point (Day1, Day2, Day 3 or Day4) by automated cell counting and growth curves were plotted. PTEN or empty vector (EV), shPTEN, shRNA to PTEN and SC, scrambled control shRNA. See also Table S1.

**Figure S2.**
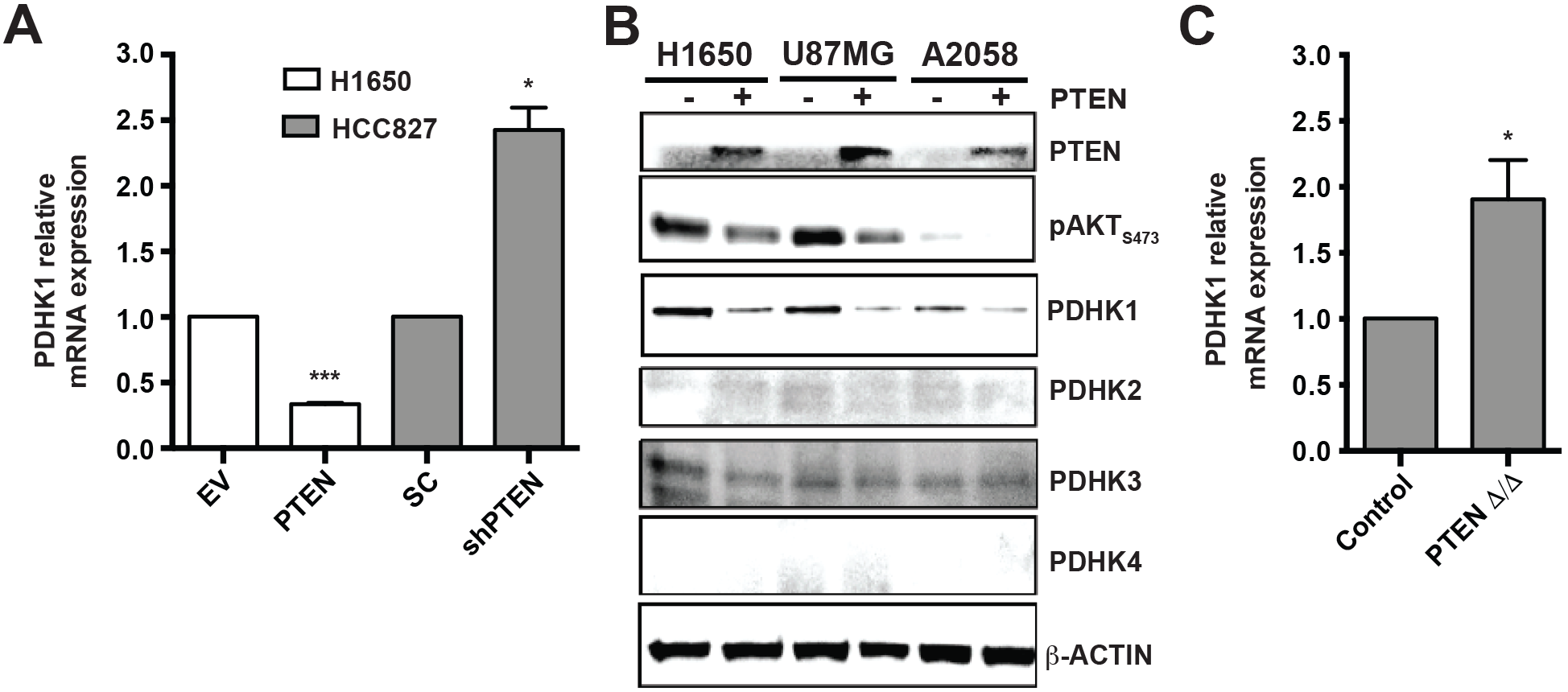
Related to Figure 1. PTEN loss upregulates energy metabolism gene *PDHK1*. (A) Quantitative real-time PCR analysis of *PDHK1* mRNA expression in PTEN-deficient H1650 cancer cell line stably expressing PTEN^WT^ or in PTEN-proficient HCC827 cancer cell line with stable *PTEN* knockdown. Data are shown as mean ± SEM (n = 3 replicates). \*\*\**p* < 0.001 compared to ‘empty vector (EV) expressing PTEN-deficient cells’ or \**p* < 0.05 compared to ‘scrambled control shRNA expressing PTEN-proficient cells’ by two-tailed unpaired t test with Welch’s correction. (B) Effects of adenovirus mediated transient PTEN re-expression in PTEN-deficient H1650, U87MG and A2058 cancer cell lines on phospho-AKT and PDHK1-4 expression by immunoblot analysis are shown. (C) Relative mRNA expression of *PDHK1* by microarray analysis in lung epithelial cells with conditional *PTEN* deletion *in vivo* (GSE47520). Data are shown as mean ± SEM (n = 3 replicates). \**p* < 0.05 compared to ‘PTEN expressing normal lung epithelial control cells’ by two-tailed unpaired t test with welch’s correction. *PDHK1* probes: 1435836_at, 1423748_at, 1434974_at and 1423747_at.

**Figure S3.**
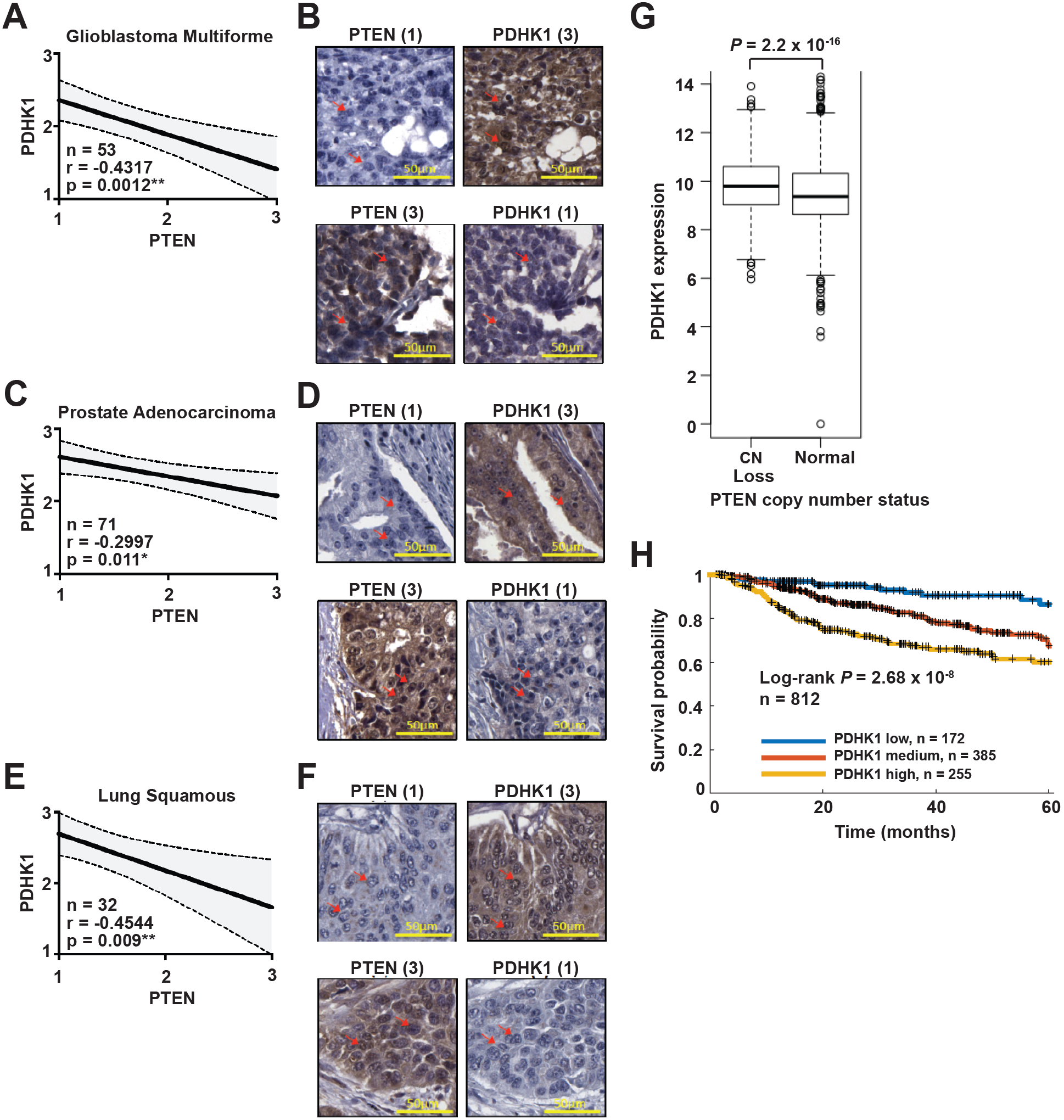
Related to Figure 1. PTEN-deficient human tumors harbor increased PDHK1: a biomarker of worse patient survival. (A, C, E) Pearson correlation analysis to assess the relationship between PTEN and PDHK1 expression in the indicated patient tumor types as assessed by immunohistochemistry staining (Methods). n, number of tumors analyzed, r, Pearson correlation coefficient where a negative value indicates inverse correlation and *p*-value, the significance of the correlation. \*\**p* < 0.01, \**p* < 0.05, as determined by two-tailed paired t-test analysis. (B, D, F) Representative IHC staining for PTEN and PDHK1 performed on tissue microarrays generated in the indicated tumor types are shown. PTEN or PDHK1 expression scored on a scale of 1 to 3, with 1 indicating ‘low’, 2 ‘intermediate’, and 3 ‘high’ expression of each protein. Scale bar, 50 μm and arrows indicate representative tumor cells scored for PTEN and PDHK1 expression. (G) Box plots indicating median (black bar), interquartile range (black box) and total range (whiskers) of normalized *PDHK1* mRNA expression levels in TCGA multi tissue pan-cancer (12 cancer types) with *PTEN* normal (n = 5995) or single or double copy number (CN) loss (n = 2845) are shown. Unfilled circles are outliers of *PDHK1* mRNA expression in these two subgroups. *P* = 2.2 × 10^−16^ ‘compared to *PTEN* normal samples’ by twotailed unpaired t test with Welch’s correction. (H) 5-year survival Kaplan-Meier analysis of PTEN CN loss patients with either high, intermediate or low level of normalized *PDHK1* mRNA expression in the cancer datasets in G.

**Figure S4.**
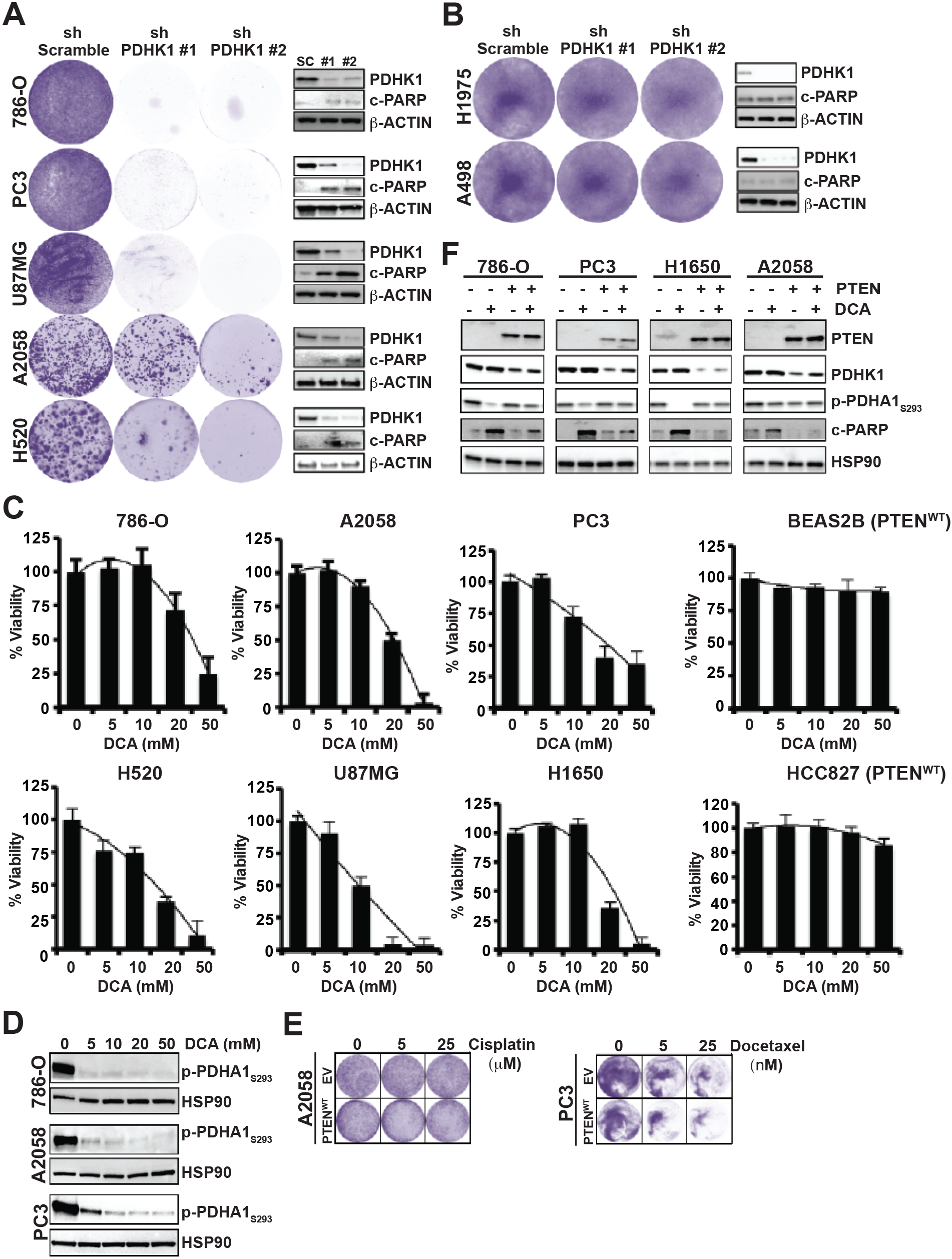
Related to Figure 2. PTEN loss or inactivation renders cells dependent on PDHK1 for survival. (A) Effects of stable *PDHK1* knockdown in various PTEN-deficient (A) or -proficient cancer cell lines (B) on cell growth by crystal violet staining assay (left) and apoptosis induction as measured by cleaved PARP levels by immunoblot analysis (right) are shown. shPDHK1#1 and shPDHK1#2, shRNAs to PDHK1 and shScramble, scrambled control shRNA. (C) Effects of pharmacologic inhibition of PDHK1 in PTEN-deficient or -proficient cancer or normal cell lines with DCA (dose response: 0, 5, 10, 20 and 50 mM) treatment or 72 hours on cell viability by CellTiter-Glo luminescent assay are shown. Data are shown as mean ± SD. (D) Western blots showing phospho-PDHA1 expression in PTEN-deficient cancer cell lines in response to PDHK1 inhibition by DCA (dose response: 0, 5, 10, 20, 50 mM) treatment. (E) Effects of chemotherapeutic agents used at indicated concentrations on cell growth of PTEN-deficient cancer cell lines stably expressing PTEN^WT^ or empty vector (EV) by crystal violet staining assay are shown. (F) Effects of pharmacologic inhibition of PDHK1 with 25 mM DCA in PTEN-deficient cancer cell lines with or without stable PTEN re-expression on pyruvate dehydrogenase complex (PDC) activation and apoptosis induction as measured by phospho-PDHA1 and cleaved PARP levels, respectively, by western blot are shown.

**Figure S5.**
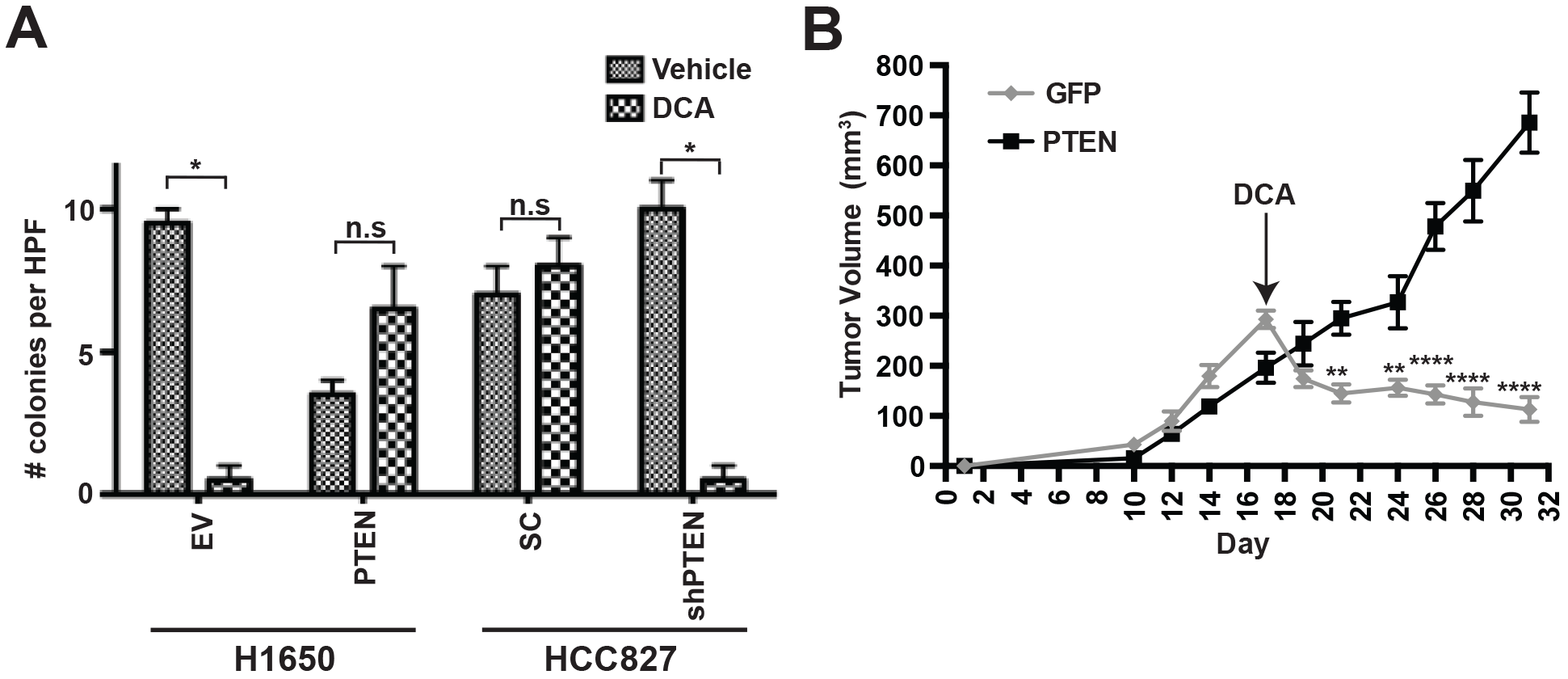
Related to Figure 2. PTEN deficiency combined with PDHK1 inhibition suppresses colony formation *in-vitro* and tumor xenograft growth *in-vivo*. (A) Effects of pharmacologic inhibition of PDHK1 with DCA in PTEN-deficient H1650 cancer cell line stably expressing PTEN or PTEN-proficient HCC827 cancer cell line with stable *PTEN* knockdown on colony formation by *in-vitro* colony formation assay are shown as measured by the number of colonies counted per high-power field (HPF: 400 x magnification). Data are shown as mean ± SEM (n = 3 replicates). \**p* < 0.05; n.s, not significant compared to ‘vehicle treated cells’ by Student’s t test. (B) Effects of pharmacologic inhibition of PDHK1 with DCA in A2058-PTEN or GFP melanoma xenografts on tumor growth *in-vivo*. Data are shown as mean ± SEM, n = 6 tumors/ group. \*\**p* < 0.01, \*\*\*\**p* < 0.0001 compared to ‘PTEN deficient GFP expressing A2058 cells’ by two-way ANOVA Sidak’s multiple comparisons test.

**Figure S6.**
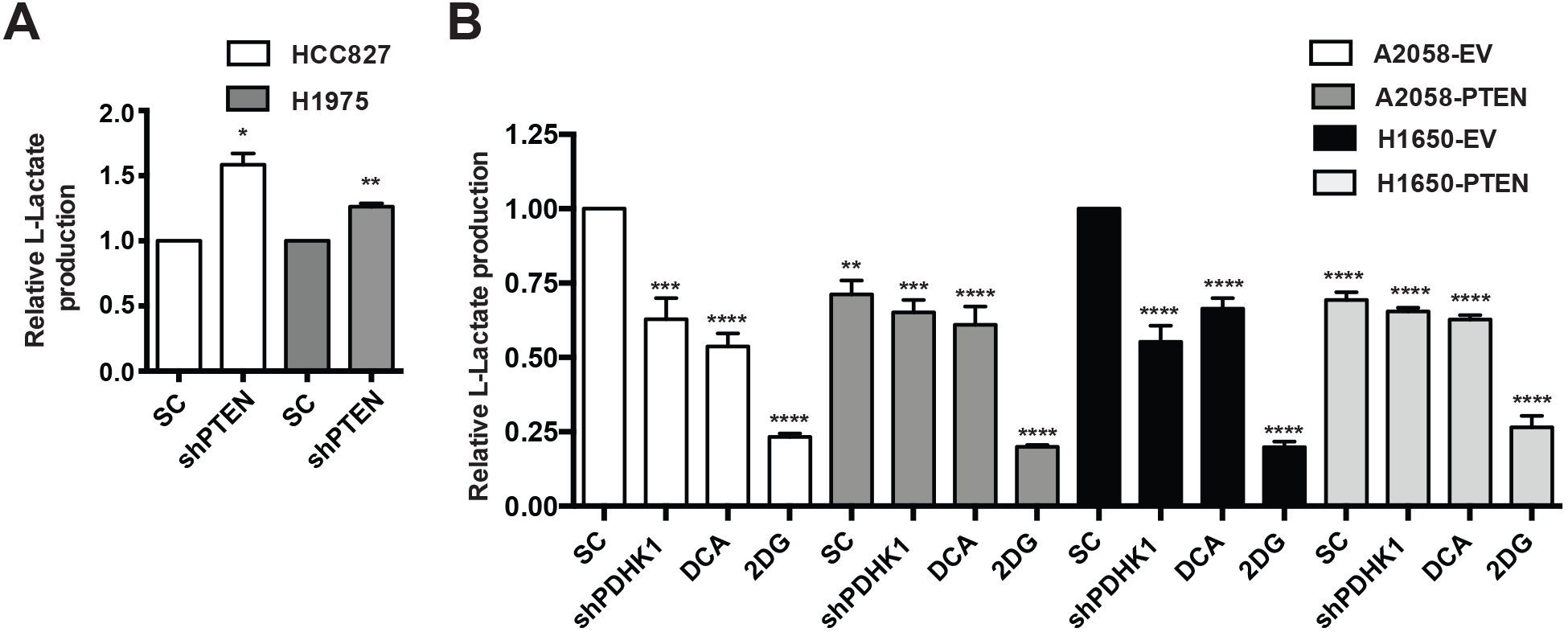
Related to Figure 2. PTEN loss promotes aerobic glycolysis via PDHK1. (A) Effects of stable *PTEN* knockdown in PTEN-proficient cancer cell lines on L-lactate production by measuring extracellular lactate in an in-vitro coupled fluorescent assay (Methods) are shown. shPTEN, shRNA to PTEN and SC, scrambled control shRNA. Data are shown as mean ± SEM (n = 3 replicates). \**p* < 0.05; \*\**p* < 0.01compared to ‘scrambled control shRNA expressing PTEN-deficient cells’ by two-tailed unpaired t test with Welch’s correction. (B) Effects of stable *PDHK1* knockdown or PDHK1 inhibition with 10 mM DCA in PTEN-deficient cancer cells stably expressing PTEN or empty vector on L-lactate production by measuring extracellular lactate in an in-vitro coupled fluorescent assay are shown. 2DG (2-deoxyglucose) was used as a +ve control to suppress lactate production independently. shPDHKI, shRNA to PDHK1 and SC, scrambled control shRNA. Data are shown as mean ± SEM (n = 3 replicates). \*\*\*\**p* < 0.0001; \*\*\**p* < 0.001; \*\**p* < 0.01 compared to ‘scrambled control shRNA expressing PTEN-deficient cells’ by Dunnett‘s multiple comparisons one-way ANOVA test.

**Figure S7.**
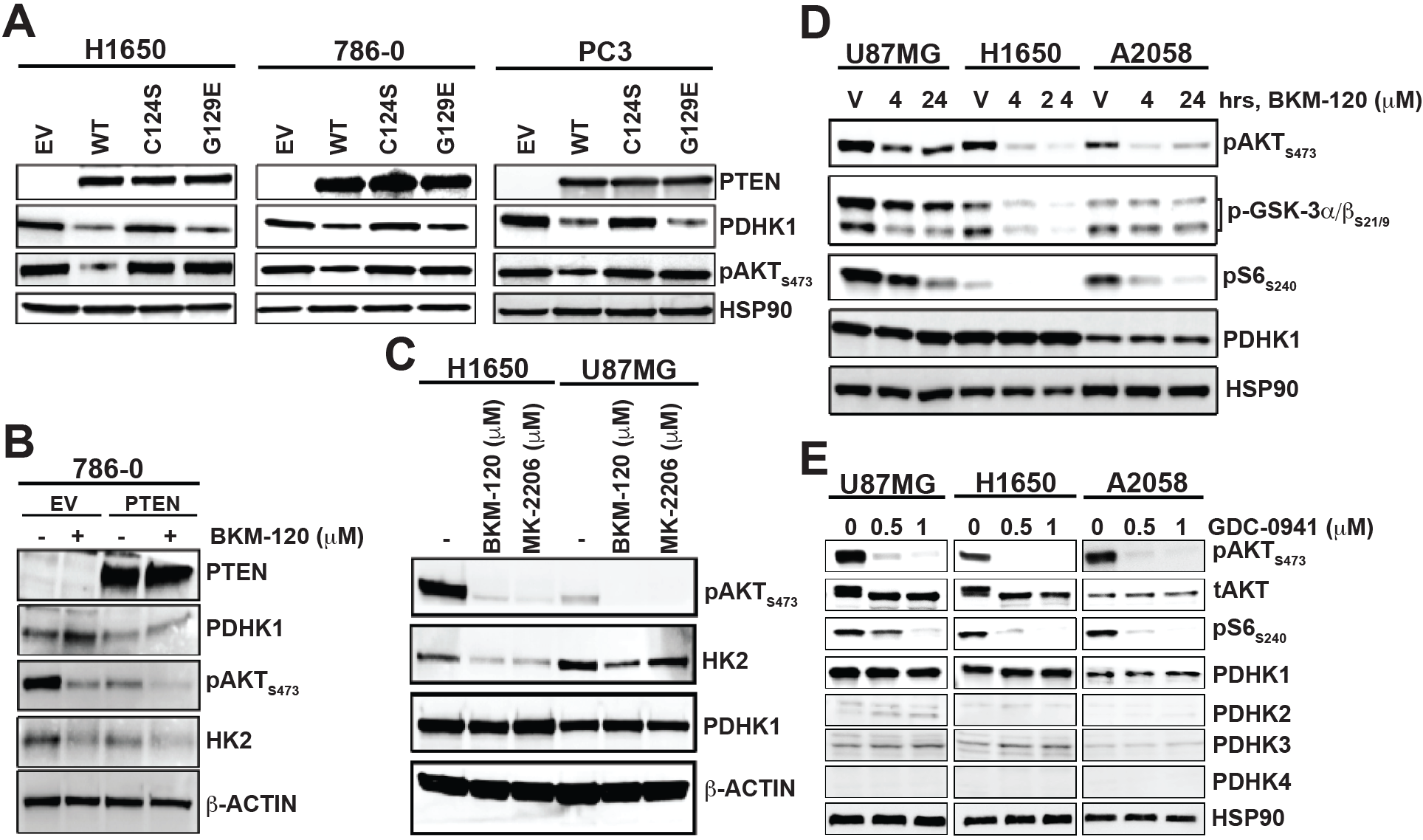
Related to Figure 3. PTEN regulates PDHK1 via its protein, and not lipid, phosphatase activity in a PI3K-AKT independent manner. (A) Western blots showing PTEN, PDHK1 and phospho-AKT expression in PTEN-deficient cancer cell lines stably expressing PTEN^WT^ or PTEN^G129E^ or PTEN^C124S^ or empty vector (EV). (B) Effects of PI3K inhibition in BKM-120 or vehicle treated PTEN-deficient 786-O cancer cells stably expressing PTEN^WT^ or empty vector (EV) for 24 hours on PDHK1, phospho-AKT and HK2 expression by immunoblotting analysis are shown. (C) Effects of PI3K or AKT inhibition in PTEN-deficient cancer cell lines treated for 24 hours with BKM-120 or MK-2206 on phospho-AKT, HK2 and PDHK1 expression by immunoblotting analysis are shown. (D) Effects of PI3K inhibition in PTEN-deficient cancer cell lines treated for 4-24 hours with BKM120 on phospho-AKT, phospho-GSK3, phospho-S6 and PDHK1 expression by immunoblotting analysis are shown. V: vehicle (DMSO) treatment. (E) Effects of PI3K inhibition in PTEN-deficient cancer cell lines treated with GDC-0941 (0, 0.5, 1 μM) for 24 hours on phospho-AKT, total AKT, phospho-S6 and PDHK1-4 expression by immunoblotting analysis are shown. PI3K or AKT inhibitors were used at 1 μM final concentration for the indicated times in B-D.

**Figure S8.**
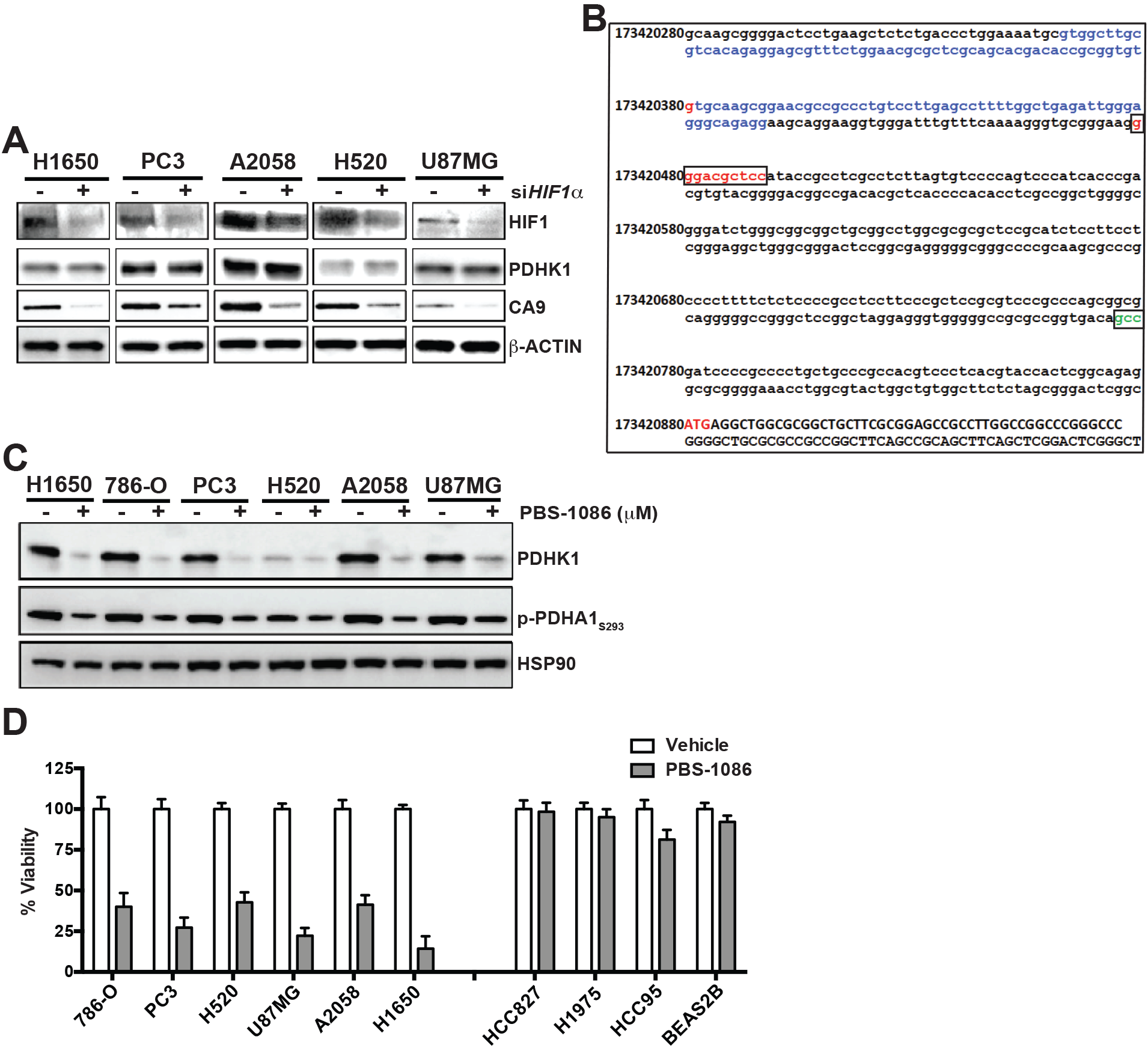
Related to Figure 4. PTEN regulates PDHK1 via NFκB and PTEN deficiency combined with NFκB inhibition decreases cell viability. (A) Effects of transient *HIF1a* knockdown in PTEN-deficient cancer cell lines on HIFlα, PDHK1 and CA9 expression by immunoblotting are shown. siHIFlα, HIFlα specific small interfering RNA and sc (-), scrambled control siRNA. (B) PDHK1 promoter sequence analysis. The canonical NFκB binding site (GGGACGCTCC, highlighted in red and boxed) is located between nucleotide positions 173420479 and 173420488 in chromosome 2, ~300 bp upstream of the transcription start site (TSS, highlighted in green and boxed). 118 bp region (highlighted in blue) spanning nucleotide position 173420380 (highlighted in red) ~40 bp upstream of the NFκB binding site in the *PDHK1* promoter was probed for NFκB recruitment by ChIP assay. ATG: start of *PDHK1* gene, highlighted in red. (C) Western blots showing PDHK1 and phospho-PDHA1 expression in PTEN-deficient cancer cell lines in response to 10μm PBS-1086 (NFκB or REL inhibitor) or vehicle (DMSO) treatment. (D) Effects of pharmacologic inhibition of NFκB in PTEN-deficient (left) or - proficient (right) cancer cell lines treated with 10 μM PBS-1086 (NFκB or REL inhibitor) or vehicle (DMSO) for 72 hours on cell viability by CellTiter-Glo luminescent assay are shown. Data are shown as mean ± SD.

**Figure S9.**
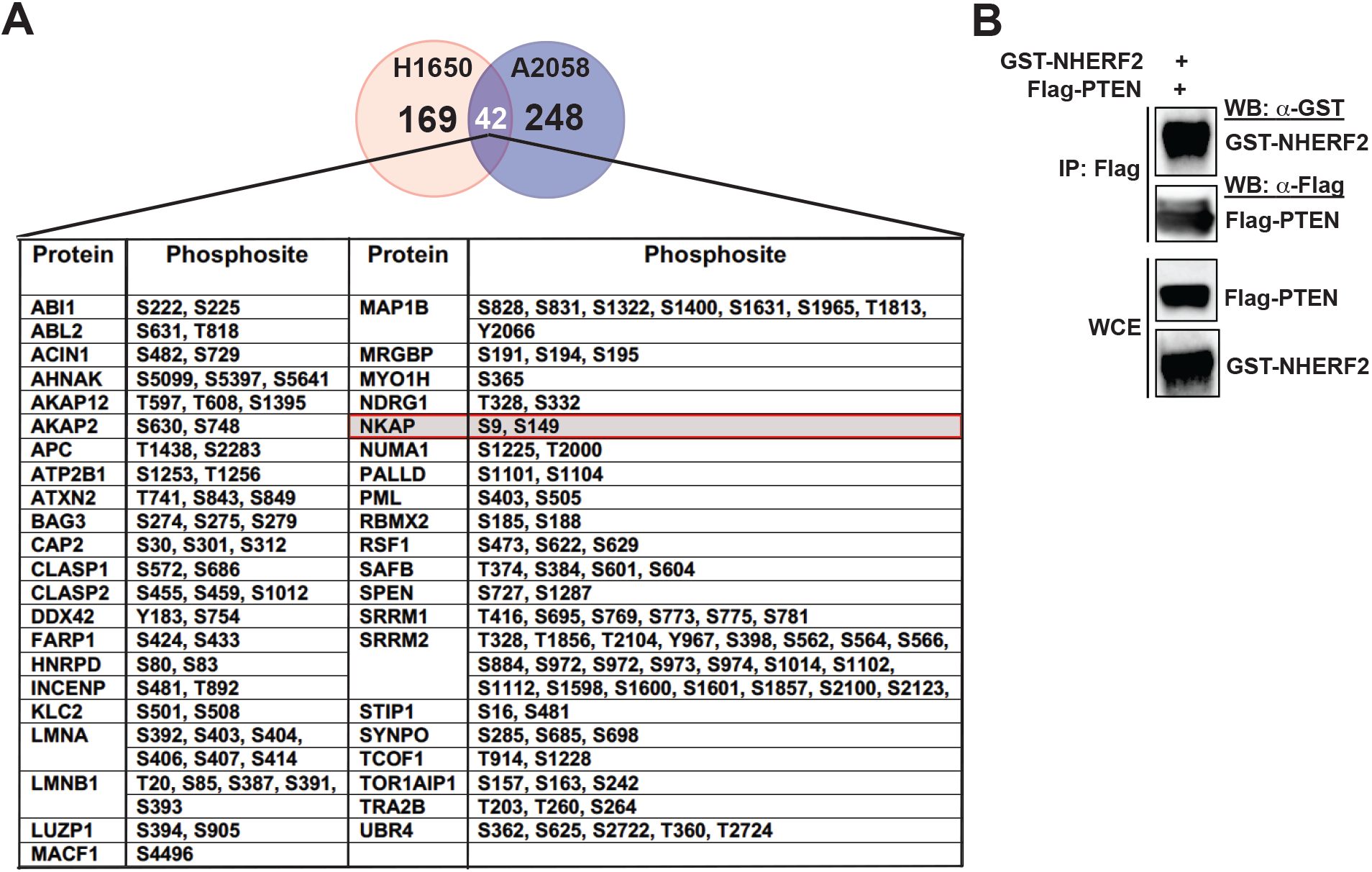
Related to Figure 5. Identification of phospho-proteins, including NKAP, that are specifically regulated by the PTEN protein phosphatase activity in cells. (A) Phospho-proteins (n=42, listed alphabetically), including NKAP (highlighted in red box), regulated specifically by the PTEN protein-phosphatase in both H1650 and A2058 PTEN-deficient cancer cell lines stably expressing either PTEN^WT^ or PTEN^G129E^ or PTEN^Y138L^ or GFP identified by an unbiased global phospho-proteomic profiling analysis (Methods) are shown. Venn diagram representation of the phospho-proteins regulated specifically by the PTEN protein phosphatase in either H1650 (n = 169) or A2058 (n = 248) or both (n = 42) cell lines. (B) Co-immunoprecipitation of NHERF2 with PTEN upon over-expression of both GST-NHERF2 and FLAG-PTEN in 293T cells followed by IP-FLAG is shown.

**Table S1.** Related to Figure 1. Genetic and expression status of PTEN in cell lines. The genetic status of PTEN in cancer and normal cell lines was determined from the Cancer Cell Line Encyclopedia (CCLE). Red (-) indicates PTEN-deficient and blue (+) indicates PTEN-expressing cells.

| Cell line | Type | Origin | PTEN gene status | Expression |
| --- | --- | --- | --- | --- |
| H1650 | Lung Adenocarcinoma | Lung | HOMDEL (exon 8,9) | - |
| H1975 | Lung Adenocarcinoma | Lung | No alteration | + |
| HCC827 | Lung Adenocarcinoma | Lung | No alteration | + |
| HCC95 | Lung Squamous Carcinoma | Lung | No alteration | + |
| PC3 | Prostate Adenocarcinoma | Prostate | R55fs*1 | - |
| 786-O | Renal Clear Cell Carcinoma | Kidney | Q149* (STOP) | - |
| H520 | Lung Squamous Carcinoma | Lung | No alteration | - |
| A2058 | Skin Cutaneous Melanoma | Skin | L112Q | - |
| U87MG | Glioblastoma Multiforme | Brain | L70_splice | - |
| 293T | Human Embryonic Kidney | Kidney | No alteration | + |
| BEAS2B | Bronchial Epithelial Normal Cells | Bronchus | No alteration | + |

**Table S2.**
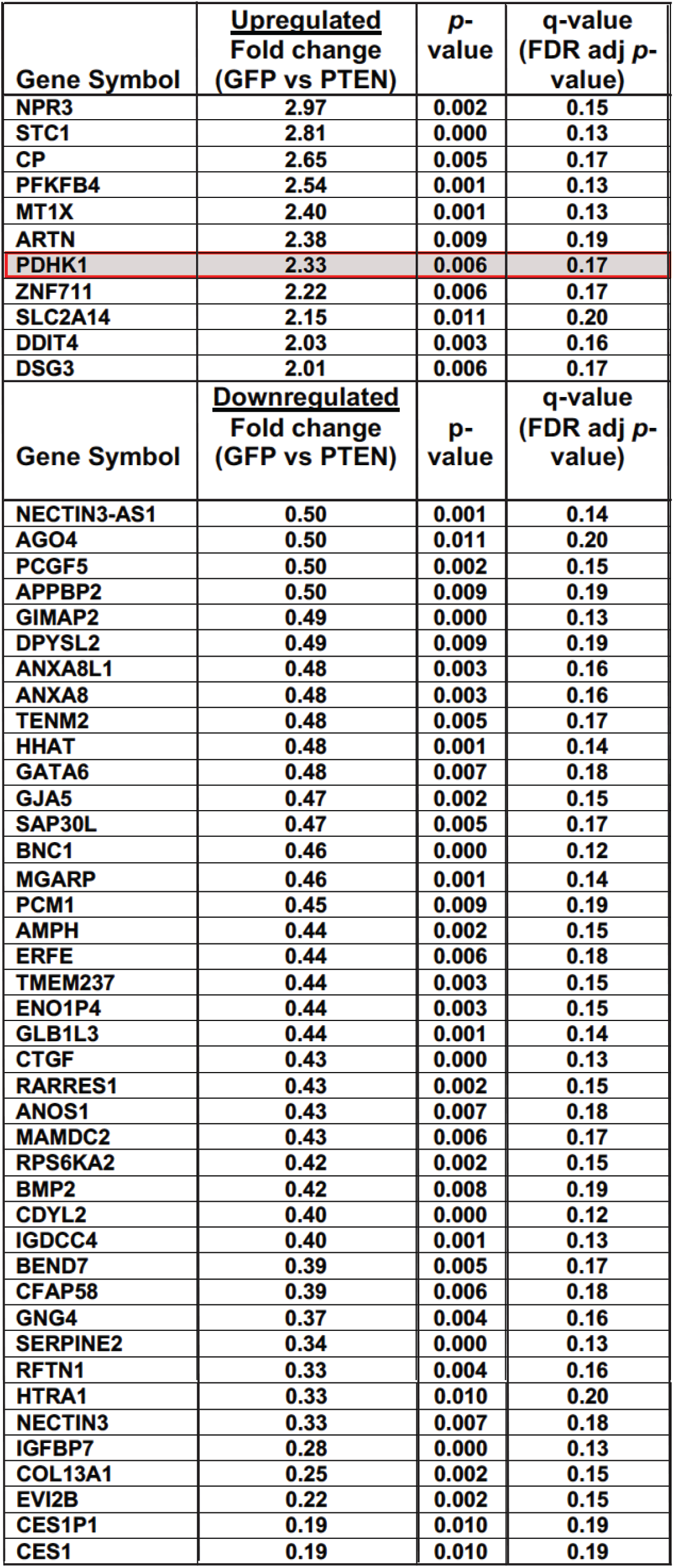
Related to Figure 1. List of genes significantly differentially expressed in response to PTEN status. Genes upregulated (including *PDHK1*, highlighted in red box) or downregulated in PTEN-deficient H1650 cells are listed. 52 genes were significantly differentially expressed between stable PTEN and GFP expressing H1650 cells by microarray analysis (fold change > 2, multiple t-test \**p* < 0.05, FDR adjusted p-value or *q* < 0.2).

**Table S3.** Related to Figure 4. List of transcription factors with consensus binding sites in the PDHK1 promoter, including NFκB. Human *PDK1* (PDHK1) promoter sequence (−499 bp from TSS) was retrieved from Eukaryotic Promoter Database (https://epd.vital-it.ch/index.php) and analyzed for transcription factor binding sites using TFBIND tool (http://tfbind.hgc.jp/) based on position weight matrix algorithm (TRANSFAC R.3.4) (Tsunoda and Takagi, 1999). Eighty-two different transcription factors, including NFκB (highlighted in red), with high TF binding score (> 0.8) were identified and are listed.

|  |  |  |  |
| --- | --- | --- | --- |
| AHR | E2 | HSF2 | PAX2 |
| AHRARNT | E2F | IK1 | PAX5 |
| AML1 | E47 | IK2 | PAX6 |
| AP1 | E4BP4 | IK3 | R |
| AP2 | EGR1 | LMO2COM | RREB1 |
| AP4 | EGR2 | LYF1 | SEF1 |
| ARP1 | EGR3 | MYOD | SP1 |
| ATF | ELK1 | MYOGNF1 | SREBP1 |
| CAAT | ER | MZF1 | SRY |
| CAP | GATA1 | NF1 | STAF |
| CDPCR3 | GATA2 | NFκB | STAT |
| CEBP | GATA3 | NFY | T3R |
| CEBPB | GC | NGFIC | TATA |
| CETS1P54 | GFI1 | NMYC | TAXCREB |
| CMYB | GR | NRF2 | USF |
| COMP1 | GRE | NRSF | VBP |
| CP2 | HEN1 | 1-Oct | VMYB |
| CREB | HLF | OLF1 | YY1 |
| CREBP1 | HNF4 | P300 | ZID |
| DELTAEF1 | HSF1 | P53 |  |

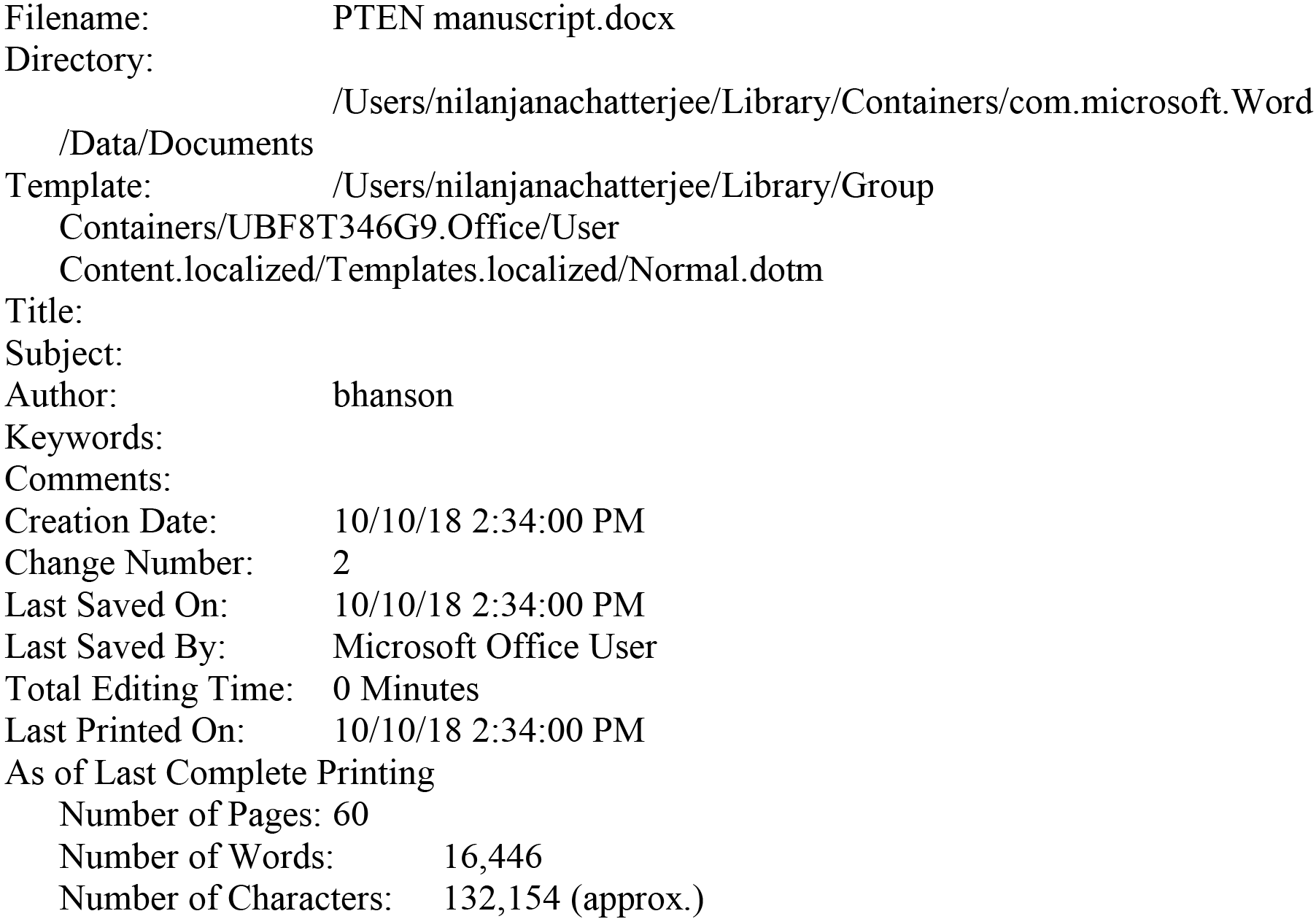

## REFERENCES

Aguissa-Toure, A. H., and Li, G. (2012). Genetic alterations of PTEN in human melanoma. Cellular and molecular life sciences : CMLS 69 1475–1491.

Aoki, Y., and Kao, P. N. (1997). Cyclosporin A-sensitive calcium signaling represses NFkappaB activation in human bronchial epithelial cells and enhances NFkappaB activation in Jurkat T-cells. Biochem Biophys Res Commun 234, 424–431.

Backman, S. A., Stambolic, V., Suzuki, A., Haight, J., Elia, A., Pretorius, J., Tsao, M. S., Shannon, P., Bolon, B., Ivy, G. O., and Mak, T. W. (2001). Deletion of Pten in mouse brain causes seizures, ataxia and defects in soma size resembling Lhermitte-Duclos disease. Nat Genet 29, 396–403.

Beltrao, P., Albanese, V., Kenner, L. R., Swaney, D. L., Burlingame, A., Villen, J., Lim, W. A., Fraser, J. S., Frydman, J., and Krogan, N. J. (2012). Systematic functional prioritization of protein posttranslational modifications. Cell 150, 413–425.

Blakely, C. M., Pazarentzos, E., Olivas, V., Asthana, S., Yan, J. J., Tan, I., Hrustanovic, G., Chan, E., Lin, L., Neel, D. S., et al. (2015). NF-kappaB-activating complex engaged in response to EGFR oncogene inhibition drives tumor cell survival and residual disease in lung cancer. Cell Rep 11, 98–110.

Burger, M. T., Pecchi, S., Wagman, A., Ni, Z. J., Knapp, M., Hendrickson, T., Atallah, G., Pfister, K., Zhang, Y., Bartulis, S., et al. (2011). Identification of NVP-BKM120 as a Potent, Selective, Orally Bioavailable Class I PI3 Kinase Inhibitor for Treating Cancer. ACS Med Chem Lett 2, 774–779.

Burgute, B. D., Peche, V. S., Steckelberg, A. L., Glockner, G., Gassen, B., Gehring, N. H., and Noegel, A. A. (2014). NKAP is a novel RS-related protein that interacts with RNA and RNA binding proteins. Nucleic Acids Res 42, 3177–3193.

Chalhoub, N., and Baker, S. J. (2009). PTEN and the PI3-kinase pathway in cancer. Annu Rev Pathol 4, 127–150.

Chandarlapaty, S., Sawai, A., Scaltriti, M., Rodrik-Outmezguine, V., Grbovic-Huezo, O., Serra, V., Majumder, P. K., Baselga, J., and Rosen, N. (2011). AKT inhibition relieves feedback suppression of receptor tyrosine kinase expression and activity. Cancer Cell 19, 58–71.

Chen, D., Li, Z., Yang, Q., Zhang, J., Zhai, Z., and Shu, H. B. (2003). Identification of a nuclear protein that promotes NF-kappaB activation. Biochem Biophys Res Commun 310, 720–724.

Chesney, J., Clark, J., Klarer, A. C., Imbert-Fernandez, Y., Lane, A. N., and Telang, S. (2014). Fructose-2,6-bisphosphate synthesis by 6-phosphofructo-2-kinase/fructose-2,6-bisphosphatase 4 (PFKFB4) is required for the glycolytic response to hypoxia and tumor growth. Oncotarget 5, 6670–6686.

Chiao, P. J., and Ling, J. (2011). Kras, Pten, NF-kappaB, and inflammation: dangerous liaisons. Cancer discovery 1, 103–105.

Choi, M., Chang, C. Y., Clough, T., Broudy, D., Killeen, T., MacLean, B., and Vitek, O. (2014). MSstats: an R package for statistical analysis of quantitative mass spectrometry-based proteomic experiments. Bioinformatics 30, 2524–2526.

Cross, D. A., Alessi, D. R., Cohen, P., Andjelkovich, M., and Hemmings, B. A. (1995). Inhibition of glycogen synthase kinase-3 by insulin mediated by protein kinase B. Nature 378, 785–789.

Dan, H. C., Cooper, M. J., Cogswell, P. C., Duncan, J. A., Ting, J. P., and Baldwin, A. S. (2008). Akt-dependent regulation of NF-{kappa}B is controlled by mTOR and Raptor in association with IKK. Genes & development 22, 1490–1500.

Davidson, L., Maccario, H., Perera, N. M., Yang, X., Spinelli, L., Tibarewal, P., Glancy, B., Gray, A., Weijer, C. J., Downes, C. P., and Leslie, N. R. (2010). Suppression of cellular proliferation and invasion by the concerted lipid and protein phosphatase activities of PTEN. Oncogene 29, 687–697.

Dey, N., Crosswell, H. E., De, P., Parsons, R., Peng, Q., Su, J. D., and Durden, D. L. (2008). The protein phosphatase activity of PTEN regulates SRC family kinases and controls glioma migration. Cancer Res 68, 1862–1871.

Fabre, C., Mimura, N., Bobb, K., Kong, S. Y., Gorgun, G., Cirstea, D., Hu, Y., Minami, J., Ohguchi, H., Zhang, J., et al. (2012). Dual inhibition of canonical and noncanonical NF-kappaB pathways demonstrates significant antitumor activities in multiple myeloma. Clinical cancer research : an official journal of the American Association for Cancer Research 18, 4669–4681.

Figueiredo, A. L., Maczkowiak, F., Borday, C., Pla, P., Sittewelle, M., Pegoraro, C., and Monsoro-Burq, A. H. (2017). PFKFB4 control of AKT signaling is essential for premigratory and migratory neural crest formation. Development 144, 4183–4194.

Folkes, A. J., Ahmadi, K., Alderton, W. K., Alix, S., Baker, S. J., Box, G., Chuckowree, I. S., Clarke, P. A., Depledge, P., Eccles, S. A, et al. (2008). The identification of 2-(1H-indazol-4-yl)-6-(4-methanesulfonyl-piperazin-1-ylmethyl)-4-morpholin-4-yl-t hieno[3,2-d]pyrimidine (GDC-0941) as a potent, selective, orally bioavailable inhibitor of class I PI3 kinase for the treatment of cancer. J Med Chem 51, 5522–5532.

Freeman, D. J., Li, A. G., Wei, G., Li, H. H., Kertesz, N., Lesche, R., Whale, A. D., Martinez-Diaz, H., Rozengurt, N., Cardiff, R. D., et al. (2003). PTEN tumor suppressor regulates p53 protein levels and activity through phosphatase-dependent and -independent mechanisms. Cancer Cell 3, 117–130.

Fulton, D., Gratton, J. P., McCabe, T. J., Fontana, J., Fujio, Y., Walsh, K., Franke, T. F., Papapetropoulos, A., and Sessa, W. C. (1999). Regulation of endothelium-derived nitric oxide production by the protein kinase Akt. Nature 399, 597–601.

Ghosh, S., Varela, L., Sood, A., Park, B. H., and Lotan, T. L. (2013). mTOR signaling feedback modulates mammary epithelial differentiation and restrains invasion downstream of PTEN loss. Cancer Res 73, 5218–5231.

Gildea, J. J., Herlevsen, M., Harding, M. A., Gulding, K. M., Moskaluk, C. A., Frierson, H. F., and Theodorescu, D. (2004). PTEN can inhibit in vitro organotypic and in vivo orthotopic invasion of human bladder cancer cells even in the absence of its lipid phosphatase activity. Oncogene 23, 6788–6797.

Gilmore, T. D. (2006). Introduction to NF-kappaB: players, pathways, perspectives. Oncogene 25, 6680–6684.

Gottlob, K., Majewski, N., Kennedy, S., Kandel, E., Robey, R. B., and Hay, N. (2001). Inhibition of early apoptotic events by Akt/PKB is dependent on the first committed step of glycolysis and mitochondrial hexokinase. Genes & development 15, 1406–1418.

Grassian, A. R., Metallo, C. M., Coloff, J. L., Stephanopoulos, G., and Brugge, J. S. (2011). Erk regulation of pyruvate dehydrogenase flux through PDK4 modulates cell proliferation. Genes Dev 25, 1716–1733.

Gu, T., Zhang, Z., Wang, J., Guo, J., Shen, W. H., and Yin, Y. (2011). CREB is a novel nuclear target of PTEN phosphatase. Cancer Res 71, 2821–2825.

Gustin, J. A., Maehama, T., Dixon, J. E., and Donner, D. B. (2001). The PTEN tumor suppressor protein inhibits tumor necrosis factor-induced nuclear factor kappa B activity. J Biol Chem 276, 27740–27744.

Hirai, H., Sootome, H., Nakatsuru, Y., Miyama, K., Taguchi, S., Tsujioka, K., Ueno, Y., Hatch, H., Majumder, P.K., Pan, B. S., and Kotani, H. (2010). MK-2206, an allosteric Akt inhibitor, enhances antitumor efficacy by standard chemotherapeutic agents or molecular targeted drugs in vitro and in vivo. Mol Cancer Ther 9, 1956–1967.

Hlobilkova, A., Guldberg, P., Thullberg, M., Zeuthen, J., Lukas, J., and Bartek, J. (2000). Cell cycle arrest by the PTEN tumor suppressor is target cell specific and may require protein phosphatase activity. Exp Cell Res 256, 571–577.

Hollander, M. C., Blumenthal, G. M., and Dennis, P. A. (2011). PTEN loss in the continuum of common cancers, rare syndromes and mouse models. Nat Rev Cancer 11, 289–301.

Houddane, A., Bultot, L., Novellasdemunt, L., Johanns, M., Gueuning, M. A., Vertommen, D., Coulie, P. G., Bartrons, R., Hue, L., and Rider, M. H. (2017). Role of Akt/PKB and PFKFB isoenzymes in the control of glycolysis, cell proliferation and protein synthesis in mitogen-stimulated thymocytes. Cell Signal 34, 23–37.

Jager, S., Cimermancic, P., Gulbahce, N., Johnson, J. R., McGovern, K. E., Clarke, S. C., Shales, M., Mercenne, G., Pache, L., Li, K., et al. (2012). Global landscape of HIV-human protein complexes. Nature 481, 365–370.

Jang, M., Kim, S. S., and Lee, J. (2013). Cancer cell metabolism: implications for therapeutic targets. Experimental & molecular medicine 45, e45.

Kato, M., Li, J., Chuang, J. L., and Chuang, D. T. (2007). Distinct structural mechanisms for inhibition of pyruvate dehydrogenase kinase isoforms by AZD7545, dichloroacetate, and radicicol. Structure 15, 992–1004.

Kim, J. W., Tchernyshyov, I., Semenza, G. L., and Dang, C. V. (2006). HIF-1-mediated expression of pyruvate dehydrogenase kinase: a metabolic switch required for cellular adaptation to hypoxia. Cell metabolism 3, 177–185.

Kinoshita, E., Kinoshita-Kikuta, E., and Koike, T. (2009). Separation and detection of large phosphoproteins using Phos-tag SDS-PAGE. Nat Protoc 4, 1513–1521.

Korotchkina, L. G., and Patel, M. S. (2001). Probing the mechanism of inactivation of human pyruvate dehydrogenase by phosphorylation of three sites. J Biol Chem 276, 5731–5738.

Koul, D. (2008). PTEN signaling pathways in glioblastoma. Cancer biology & therapy 7, 1321–1325.

Koul, D., Yao, Y., Abbruzzese, J. L., Yung, W. K., and Reddy, S. A. (2001). Tumor suppressor MMAC/PTEN inhibits cytokine-induced NFkappaB activation without interfering with the IkappaB degradation pathway. J Biol Chem 276, 11402–11408.

Kwon, C. H., Luikart, B. W., Powell, C. M., Zhou, J., Matheny, S. A., Zhang, W., Li, Y., Baker, S. J., and Parada, L. F. (2006). Pten regulates neuronal arborization and social interaction in mice. Neuron 50, 377–388.

Lee, H. J., Lee, H. Y., Lee, J. H., Lee, H., Kang, G., Song, J. S., Kang, J., and Kim, J. H. (2014). Prognostic Significance of Biallelic Loss of PTEN in Clear Cell Renal Cell Carcinoma. The Journal of urology.

Lee, J. O., Yang, H., Georgescu, M. M., Di Cristofano, A., Maehama, T., Shi, Y., Dixon, J. E., Pandolfi, P., and Pavletich, N. P. (1999). Crystal structure of the PTEN tumor suppressor: implications for its phosphoinositide phosphatase activity and membrane association. Cell 99, 323–334.

Leslie, N. R., Maccario, H., Spinelli, L., and Davidson, L. (2009). The significance of PTEN’s protein phosphatase activity. Adv Enzyme Regul 49, 190–196.

Leslie, N. R., Yang, X., Downes, C. P., and Weijer, C. J. (2007). PtdIns(3,4,5)P(3)-dependent and -independent roles for PTEN in the control of cell migration. Curr Biol 17, 115–125.

Li, D. M., and Sun, H. (1997). TEP1, encoded by a candidate tumor suppressor locus, is a novel protein tyrosine phosphatase regulated by transforming growth factor beta. Cancer Res 57, 2124–2129.

Li, J., Yen, C., Liaw, D., Podsypanina, K., Bose, S., Wang, S. I., Puc, J., Miliaresis, C., Rodgers, L., McCombie, R., et al. (1997). PTEN, a putative protein tyrosine phosphatase gene mutated in human brain, breast, and prostate cancer. Science 275, 1943–1947.

Li, T., Chen, L., Cheng, J., Dai, J., Huang, Y., Zhang, J., Liu, Z., Li, A., Li, N., Wang, H., et al. (2016). SUMOylated NKAP is essential for chromosome alignment by anchoring CENP-E to kinetochores. Nat Commun 7, 12969.

Linn, T. C., Pettit, F. H., and Reed, L. J. (1969). Alpha-keto acid dehydrogenase complexes. X. Regulation of the activity of the pyruvate dehydrogenase complex from beef kidney mitochondria by phosphorylation and dephosphorylation. Proc Natl Acad Sci U S A 62, 234–241.

Maehama, T., and Dixon, J. E. (1998). The tumor suppressor, PTEN/MMAC1, dephosphorylates the lipid second messenger, phosphatidylinositol 3,4,5-trisphosphate. J Biol Chem 273, 13375–13378.

Maehama, T., and Dixon, J. E. (1999). PTEN: a tumour suppressor that functions as a phospholipid phosphatase. Trends Cell Biol 9, 125–128.

Mahimainathan, L., and Choudhury, G. G. (2004). Inactivation of platelet-derived growth factor receptor by the tumor suppressor PTEN provides a novel mechanism of action of the phosphatase. J Biol Chem 279, 15258–15268.

Maier, D., Jones, G., Li, X., Schonthal, A. H., Gratzl, O., Van Meir, E. G., and Merlo, A. (1999). The PTEN lipid phosphatase domain is not required to inhibit invasion of glioma cells. Cancer Res 59, 5479–5482.

Martone, R., Euskirchen, G., Bertone, P., Hartman, S., Royce, T. E., Luscombe, N. M., Rinn, J. L., Nelson, F. K., Miller, P., Gerstein, M., et al. (2003). Distribution of NF-kappaB-binding sites across human chromosome 22. Proc Natl Acad Sci U S A 100, 12247–12252.

Mayo, M. W., Madrid, L. V., Westerheide, S. D., Jones, D. R., Yuan, X. J., Baldwin, A. S., Jr., and Whang, Y. E. (2002). PTEN blocks tumor necrosis factor-induced NF-kappa B-dependent transcription by inhibiting the transactivation potential of the p65 subunit. J Biol Chem 277, 11116–11125.

Myers, M. P., Pass, I., Batty, I. H., Van der Kaay, J., Stolarov, J. P., Hemmings, B. A., Wigler, M. H., Downes, C. P., and Tonks, N. K. (1998). The lipid phosphatase activity of PTEN is critical for its tumor supressor function. Proc Natl Acad Sci U S A 95, 13513–13518.

Myers, M. P., Stolarov, J. P., Eng, C., Li, J., Wang, S. I., Wigler, M. H., Parsons, R., and Tonks, N. K. (1997). P-TEN, the tumor suppressor from human chromosome 10q23, is a dual-specificity phosphatase. Proc Natl Acad Sci U S A 94, 9052–9057.

Neshat, M. S., Mellinghoff, I. K., Tran, C., Stiles, B., Thomas, G., Petersen, R., Frost, P., Gibbons, J. J., Wu, H., and Sawyers, C. L. (2001). Enhanced sensitivity of PTEN-deficient tumors to inhibition of FRAP/mTOR. Proc Natl Acad Sci U S A 98, 10314–10319.

Ory, D. S., Neugeboren, B. A., and Mulligan, R. C. (1996). A stable human-derived packaging cell line for production of high titer retrovirus/vesicular stomatitis virus G pseudotypes. Proc Natl Acad Sci U S A 93, 11400–11406.

Ozers, M. S., Ervin, K. M., Steffen, C. L., Fronczak, J. A., Lebakken, C. S., Carnahan, K. A., Lowery, R. G., and Burke, T. J. (2005). Analysis of ligand-dependent recruitment of coactivator peptides to estrogen receptor using fluorescence polarization. Mol Endocrinol 19, 25–34.

Pajerowski, A. G., Nguyen, C., Aghajanian, H., Shapiro, M. J., and Shapiro, V. S. (2009). NKAP is a transcriptional repressor of notch signaling and is required for T cell development. Immunity 30, 696–707.

Patel, M. S., Nemeria, N. S., Furey, W., and Jordan, F. (2014). The pyruvate dehydrogenase complexes: structure-based function and regulation. J Biol Chem 289, 16615–16623.

Patel, M. S., and Roche, T. E. (1990). Molecular biology and biochemistry of pyruvate dehydrogenase complexes. FASEB J 4, 3224–3233.

Pathak, R. R., Grover, A., Malaney, P., Quarni, W., Pandit, A., Allen-Gipson, D., and Dave, V. (2013). Loss of phosphatase and tensin homolog (PTEN) induces leptin-mediated leptin gene expression: feed-forward loop operating in the lung. J Biol Chem 288, 29821–29835.

Planchon, S. M., Waite, K. A., and Eng, C. (2008). The nuclear affairs of PTEN. J Cell Sci 121, 249–253.

Popov, K. M., Hawes, J. W., and Harris, R. A. (1997). Mitochondrial alpha-ketoacid dehydrogenase kinases: a new family of protein kinases. Adv Second Messenger Phosphoprotein Res 31, 105–111.

Pourmand, G., Ziaee, A. A., Abedi, A. R., Mehrsai, A., Alavi, H. A., Ahmadi, A., and Saadati, H. R. (2007). Role of PTEN gene in progression of prostate cancer. Urology journal 4, 95–100.

Raftopoulou, M., Etienne-Manneville, S., Self, A., Nicholls, S., and Hall, A. (2004). Regulation of cell migration by the C2 domain of the tumor suppressor PTEN. Science 303, 1179–1181.

Rodon, J., Dienstmann, R., Serra, V., and Tabernero, J. (2013). Development of PI3K inhibitors: lessons learned from early clinical trials. Nat Rev Clin Oncol 10, 143–153.

Rodriguez-Escudero, I., Oliver, M. D., Andres-Pons, A., Molina, M., Cid, V. J., and Pulido, R. (2011). A comprehensive functional analysis of PTEN mutations: implications in tumor-and autism-related syndromes. Human molecular genetics 20, 4132–4142.

Schulze, A., and Downward, J. (2011). Flicking the Warburg switch-tyrosine phosphorylation of pyruvate dehydrogenase kinase regulates mitochondrial activity in cancer cells. Molecular cell 44, 846–848.

Schulze, A., and Harris, A. L. (2012). How cancer metabolism is tuned for proliferation and vulnerable to disruption. Nature 491, 364–373.

Semenza, G. L. (2008). Tumor metabolism: cancer cells give and take lactate. J Clin Invest 118, 3835–3837.

Shen, W. H., Balajee, A. S., Wang, J., Wu, H., Eng, C., Pandolfi, P. P., and Yin, Y. (2007). Essential role for nuclear PTEN in maintaining chromosomal integrity. Cell 128, 157–170.

Shi, Y., Wang, J., Chandarlapaty, S., Cross, J., Thompson, C., Rosen, N., and Jiang, X. (2014). PTEN is a protein tyrosine phosphatase for IRS1. Nat Struct Mol Biol 21, 522–527.

Shinde, S. R., and Maddika, S. (2016). PTEN modulates EGFR late endocytic trafficking and degradation by dephosphorylating Rab7. Nat Commun 7, 10689.

Siggers, T., Duyzend, M. H., Reddy, J., Khan, S., and Bulyk, M. L. (2011). Non-DNA-binding cofactors enhance DNA-binding specificity of a transcriptional regulatory complex. Mol Syst Biol 7, 555.

Slattery, M., Riley, T., Liu, P., Abe, N., Gomez-Alcala, P., Dror, I., Zhou, T., Rohs, R., Honig, B., Bussemaker, H. J., and Mann, R. S. (2011). Cofactor binding evokes latent differences in DNA binding specificity between Hox proteins. Cell 147, 1270–1282.

Song, M. S., Salmena, L., and Pandolfi, P. P. (2012). The functions and regulation of the PTEN tumour suppressor. Nat Rev Mol Cell Biol 13, 283–296.

Sos, M. L., Koker, M., Weir, B. A., Heynck, S., Rabinovsky, R., Zander, T., Seeger, J. M., Weiss, J., Fischer, F., Frommolt, P., et al. (2009). PTEN loss contributes to erlotinib resistance in EGFR-mutant lung cancer by activation of Akt and EGFR. Cancer research 69, 3256–3261.

St-Denis, N., Gupta, G. D., Lin, Z. Y., Gonzalez-Badillo, B., Veri, A. O., Knight, J. D. R., Rajendran, D., Couzens, A. L., Currie, K. W., Tkach, J. M., et al. (2016). Phenotypic and Interaction Profiling of the Human Phosphatases Identifies Diverse Mitotic Regulators. Cell Rep 17, 2488–2501.

Stacpoole, P. W. (1989). The pharmacology of dichloroacetate. Metabolism 38, 1124–1144.

Stambolic, V., Suzuki, A., de la Pompa, J. L., Brothers, G. M., Mirtsos, C., Sasaki, T., Ruland, J., Penninger, J. M., Siderovski, D. P., and Mak, T. W. (1998). Negative regulation of PKB/Akt-dependent cell survival by the tumor suppressor PTEN. Cell 95, 29–39.

Steck, P. A., Pershouse, M. A., Jasser, S. A., Yung, W. K., Lin, H., Ligon, A. H., Langford, L. A., Baumgard, M. L., Hattier, T., Davis, T., et al. (1997). Identification of a candidate tumour suppressor gene, MMAC1, at chromosome 10q23.3 that is mutated in multiple advanced cancers. Nat Genet 15, 356–362.

Sun, C., Wang, L., Huang, S., Heynen, G. J., Prahallad, A., Robert, C., Haanen, J., Blank, C., Wesseling, J., Willems, S. M., et al. (2014). Reversible and adaptive resistance to BRAF(V600E) inhibition in melanoma. Nature 508, 118–122.

Sun, H., Lesche, R., Li, D. M., Liliental, J., Zhang, H., Gao, J., Gavrilova, N., Mueller, B., Liu, X., and Wu, H. (1999). PTEN modulates cell cycle progression and cell survival by regulating phosphatidylinositol 3,4,5,-trisphosphate and Akt/protein kinase B signaling pathway. Proc Natl Acad Sci U S A 96, 6199–6204.

Suzuki, K., Bose, P., Leong-Quong, R. Y., Fujita, D. J., and Riabowol, K. (2010). REAP: A two minute cell fractionation method. BMC Res Notes 3, 294.

Takahashi, Y., Morales, F. C., Kreimann, E. L., and Georgescu, M. M. (2006). PTEN tumor suppressor associates with NHERF proteins to attenuate PDGF receptor signaling. EMBO J 25, 910–920.

Takeda, H., Takigawa, N., Ohashi, K., Minami, D., Kataoka, I., Ichihara, E., Ochi, N., Tanimoto, M., and Kiura, K. (2013). Vandetanib is effective in EGFR-mutant lung cancer cells with PTEN deficiency. Exp Cell Res 319, 417–423.

Taylor, G. S., and Dixon, J. E. (2003). PTEN and myotubularins: families of phosphoinositide phosphatases. Methods Enzymol 366, 43–56.

Teague, W. M., Pettit, F. H., Yeaman, S. J., and Reed, L. J. (1979). Function of phosphorylation sites on pyruvate dehydrogenase. Biochem Biophys Res Commun 87, 244–252.

Thapa, P., Chen, M. W., McWilliams, D. C., Belmonte, P., Constans, M., Sant’Angelo, D. B., and Shapiro, V. S. (2016). NKAP Regulates Invariant NKT Cell Proliferation and Differentiation into ROR-gammat-Expressing NKT17 Cells. J Immunol 196, 4987–4998.

Tibarewal, P., Zilidis, G., Spinelli, L., Schurch, N., Maccario, H., Gray, A., Perera, N. M., Davidson, L., Barton, G. J., and Leslie, N. R. (2012). PTEN protein phosphatase activity correlates with control of gene expression and invasion, a tumor-suppressing phenotype, but not with AKT activity. Sci Signal 5, ra18.

Vander Heiden, M. G., Cantley, L. C., and Thompson, C. B. (2009). Understanding the Warburg effect: the metabolic requirements of cell proliferation. Science 324, 1029–1033.

Wan, F., and Lenardo, M. J. (2009). Specification of DNA binding activity of NF-kappaB proteins. Cold Spring Harb Perspect Biol 1, a000067.

Wang, L., Xiong, H., Wu, F., Zhang, Y., Wang, J., Zhao, L., Guo, X., Chang, L. J., Zhang, Y., You, M. J., et al. (2014). Hexokinase 2-mediated Warburg effect is required for PTEN-and p53-deficiency-driven prostate cancer growth. Cell Rep 8, 1461–1474.

Warburg, O. (1956). On the origin of cancer cells. Science 123, 309–314.

Weatherman, R. V., Chang, C. Y., Clegg, N. J., Carroll, D. C., Day, R. N., Baxter, J. D., McDonnell, D. P., Scanlan, T. S., and Schaufele, F. (2002). Ligand-selective interactions of ER detected in living cells by fluorescence resonance energy transfer. Mol Endocrinol 16, 487–496.

Whitehouse, S., Cooper, R. H., and Randle, P. J. (1974). Mechanism of activation of pyruvate dehydrogenase by dichloroacetate and other halogenated carboxylic acids. Biochem J 141, 761–774.

Wittenberg, A. D., Azar, S., Klochendler, A., Stolovich-Rain, M., Avraham, S., Birnbaum, L., Binder Gallimidi, A., Katz, M., Dor, Y., and Meyuhas, O. (2016). Phosphorylated Ribosomal Protein S6 Is Required for Akt-Driven Hyperplasia and Malignant Transformation, but Not for Hypertrophy, Aneuploidy and Hyperfunction of Pancreatic beta-Cells. PLoS One 11, e0149995.

Wozniak, D. J., Kajdacsy-Balla, A., Macias, V., Ball-Kell, S., Zenner, M. L., Bie, W., and Tyner, A. L. (2017). PTEN is a protein phosphatase that targets active PTK6 and inhibits PTK6 oncogenic signaling in prostate cancer. Nat Commun 8, 1508.

Wykoff, C. C., Beasley, N. J., Watson, P. H., Turner, K. J., Pastorek, J., Sibtain, A., Wilson, G. D., Turley, H., Talks, K. L., Maxwell, P. H., et al. (2000). Hypoxia-inducible expression of tumor-associated carbonic anhydrases. Cancer Res 60, 7075–7083.

You, D., Xin, J., Volk, A., Wei, W., Schmidt, R., Scurti, G., Nand, S., Breuer, E. K., Kuo, P. C., Breslin, P., et al.(2015). FAK mediates a compensatory survival signal parallel to PI3K-AKT in PTEN-null T-ALL cells. Cell Rep 10, 2055–2068.

Zhang, X. C., Piccini, A., Myers, M. P., Van Aelst, L., and Tonks, N. K. (2012). Functional analysis of the protein phosphatase activity of PTEN. Biochem J 444, 457–464.

Zhao, Y., Butler, E. B., and Tan, M. (2013). Targeting cellular metabolism to improve cancer therapeutics. Cell death & disease 4, e532.

Zhou, J., and Parada, L. F. (2012). PTEN signaling in autism spectrum disorders. Curr Opin Neurobiol 22, 873–879.

